# Community-wide interactions sustain life in geothermal spring habitats

**DOI:** 10.1101/2024.09.03.611078

**Authors:** Timothy G. Stephens, Julia Van Etten, Timothy McDermott, William Christian, Martha Chaverra, James Gurney, Yongsung Lee, Hocheol Kim, Chung Hyun Cho, Erik Chovancek, Philipp Westhoff, Antonia Otte, Trent R. Northen, Benjamin P. Bowen, Katherine B. Louie, Kerrie Barry, Igor V. Grigoriev, Thomas Mock, Shao-Lun Liu, Shin-ya Miyagishima, Masafumi Yoshinaga, Andreas P.M. Weber, Hwan Su Yoon, Debashish Bhattacharya

**Author notes:** Corresponding authors. &.

## Abstract

We investigated an alga-dominated geothermal spring community in Yellowstone National Park, USA. Our goal was to determine how cells cope with abiotic stressors during diurnal sampling that spanned over two orders of magnitude in solar irradiance. We report a community level response to toxic metal resistance and energy cycling that spans the three domains of life. Arsenic detoxification is accomplished *via* complementary gene expression by different lineages. Photosynthesis is dominated by *Cyanidioschyzon*, with the mixotroph, *Galdieria*, relegated to nighttime heterotrophy. Many key functions, including the cell cycle, are strongly regulated by diurnal light fluctuations. These results demonstrate that biotic interactions are highly structured in extreme habitats. We suggest this was also the case on the early Earth when geothermal springs were cradles of microbial life, prior to the origin of eukaryotes.

## Introduction

Prokaryotes often live in habitats that have extremes of pH, temperature, solar radiation, salt concentrations, and atmospheric pressure. These conditions were commonplace in the early Earth and may exist in exoplanets ^1–3^. One of the few microbial eukaryotes that have evolved the capacity to inhabit such extreme environments are the photosynthetic red algae, Cyanidiophyceae ^4,5^. This unicellular lineage includes well-studied genera such as *Galdieria* and *Cyanidioschyzon* ^6^ that inhabit, and often dominate, volcanic geothermal springs and acid mining sites characterized by extremes of light levels, relatively high temperature (35-63°C), and low pH (0 to 5), with high salt and toxic heavy metal concentrations ^7,8^. Cyanidiophyceae, which are descended from mesophilic red algal ancestors, have adapted to life in geothermal springs due to horizontal gene transfer (HGT), which has led to ca. 1% of their nuclear gene inventory being comprised of HGT-derived prokaryotic genes ^4^. These foreign sequences confer polyextremophily, including metal and xenobiotic resistance and detoxification, cellular oxidant reduction, carbon metabolism, amino acid metabolism, osmotic resistance, and salt tolerance ^4,9–11^. Here, we used environmental multi-omics to investigate biotic interactions in geothermal habitats at Lemonade Creek, Yellowstone National Park (YNP), USA. Our goal was to decipher the roles of Cyanidiophyceae and prokaryotes in sustaining these communities in three distinct habitats: submerged lush biofilms, immediately adjacent endolithic sites, and acidic soils bordering the creek (**Fig. 1A**). Our sampling strategy also assessed community transcriptional and metabolic responses to solar irradiance, a major energy input for *Cyanidioschyzon* and we believe, to a lesser extent for *Galdieria*, both of which are the dominant primary producers in this environment.

**Fig. 1.**
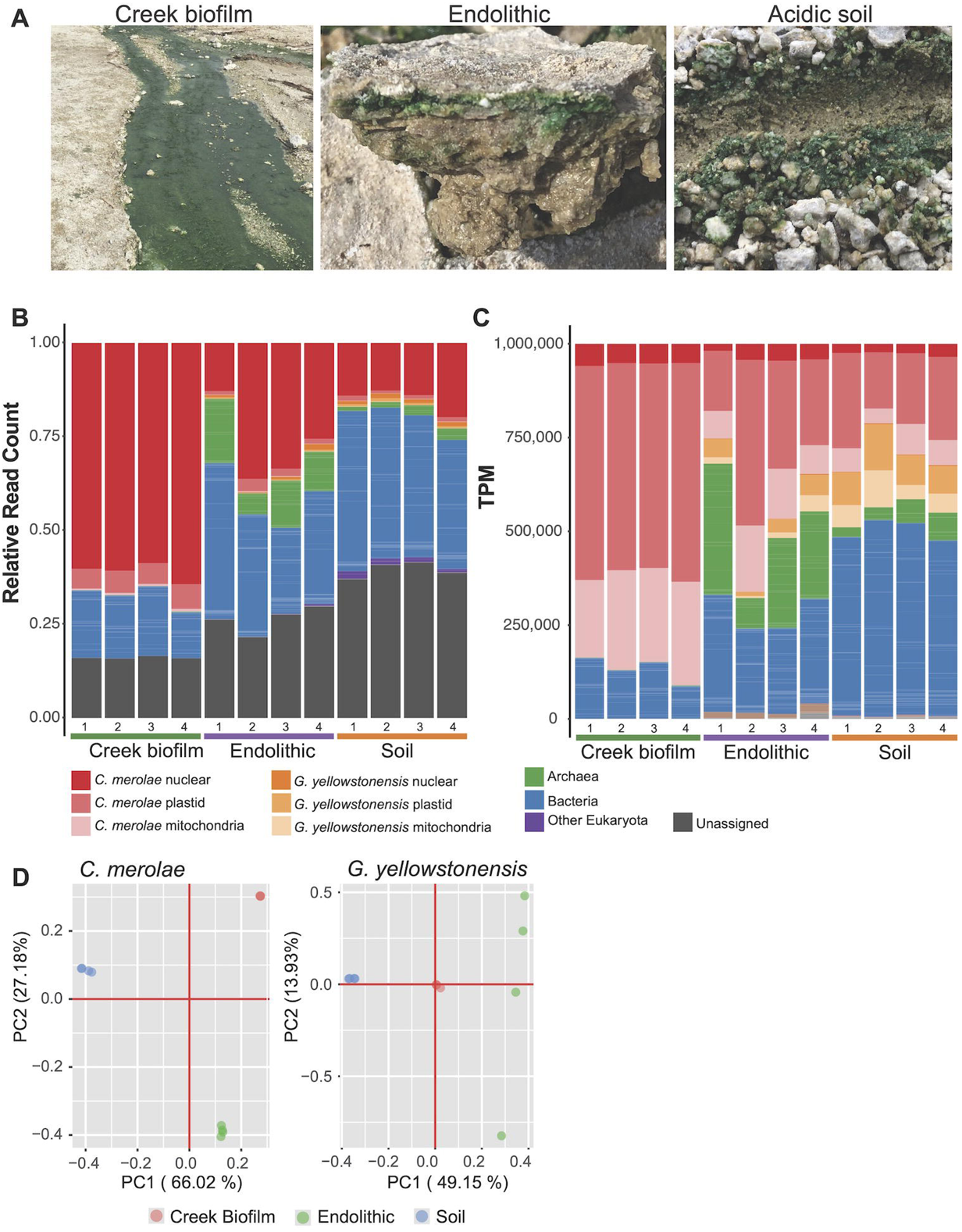
The studied YNP habitats and analysis of metagenome data. (**A**) The submerged Creek biofilm, Endolithic, and adjacent acidic Soil habitats that were sampled in Lemonade Creek, YNP. (**B**) The relative number of reads (out of the total corrected reads from a given sample) that aligned to each type of MAG (i.e., bacterial, archaeal, eukaryotic, and unassigned) and the *C. merolae* 10D and *G. yellowstonensis* 5587.1 reference nuclear and organelle genomes in the combined non-redundant dataset from the three YNP habitats. **(C)** The relative abundance, in Transcripts per Million (TPM), of each MAG. (**D**) Principal component analysis (PCA) of *C. merolae* and *G. yellowstonensis* single-nucleotide polymorphism (SNP) data from the three studied habitats at YNP. The first two principal components are shown for each species. These data demonstrate that the algal populations have a high degree of distinctness in PC1. For *G. yellowstonensis,* the Endolithic populations are dispersed largely due to lack of SNP data from the Creek biofilm population (PC1, 0; PC2, 0).

## Materials and methods

### Multi-omics sampling

Samples were collected from Lemonade Creek, a geothermal site in YNP from three distinct environments, hereafter referred to as “Creek biofilm” (dense lush green biofilms in the creek channel), “Soil” (bordering the creek channel), and “Endolithic” (biomass within rocks sampled within centimeters of the creek flow). Each environment was sampled at a single time point in quadruplicate (n=4) for Illumina short read metagenome data generation. Restrictions limiting the amount of material collected from the Endolithic environment meant that samples for metatranscriptomic (Illumina short read) and metabolomic (polar LC-MS) analysis were only collected from the Creek biofilm and Soil environments. These environments were sampled (n=4) at four time points (TP1, commencing at sunrise, 07:30; TP2, ∼ midday 12:50; TP3, early dusk, 16:50, and TP4, complete darkness, 19:25). Detailed protocols for sample collection, extraction, and data generation are presented in the **Supplemental Information**.

### MAG generation

For each sample, short read metagenome data was processed and assembled by the DOE Joint Genome Institute (JGI) following SOPs 1007.3 – LC, 1082.2, and 1077. This produced 12 metagenome assemblies (4 per environment), which had scaffolds from Cyanidiophyceae removed by a BLASTN comparison (>90% identity and >90% coverage) against the available reference genomes. This analysis also elucidated that there were two Cyanidiophyceae species present at these sites, a *Cyanidioschyzon merolae* 10D-like isolate and a *Galdieria yellowstonensis* 5587.1-like isolate [previously *G. sulphuraria* ^6^]. Prokaryotic MAGs were constructed from the Cyanidiophyceae cleaned assemblies using a comprehensive workflow, which combined multiple binning approaches, contamination assessment and cleaning, as well as per-environment reassembly and binning to aid the construction of MAGs from rare microbes (see **Supplemental Information** for details). The prokaryotic MAGs from the 12 samples and three per-environment reassemblies were merged into a final non-redundant set of MAGs using an average nucleotide identity (ANI) of 98%. This threshold is below species level, meaning that each MAG represents a district strain, and roughly correlates to the limit at which genomes are distinct when mapping short reads ^12^, which was the primary aim of our downstream analysis. Non-Cyanidiophyceae eukaryotic MAGs were constructed from the three per-environment reassemblies, although all five were identified in the Soil reassembly since it had the highest species diversity and thus produced the most contiguous assembly. Significant effort was put into ensuring these eukaryotic MAGs were free from prokaryotic contamination and had genes predicted using a taxa appropriate approach (i.e., ciliates, which we had a single MAG from, have unusual genome structure and non-standard codons which require special considerations; see **Supplemental Information**). The abundance of the non-redundant prokaryotic MAGs, the nuclear and organellar *C. merolae* 10D and *G. yellowstonensis* YNP5587_1 reference genomes, eukaryotic MAGs, and viral vOTUs (hereinafter, combined MAG dataset) assembled from the same samples in a previous analysis ^13^, were quantified across the 12 metagenome samples (encompassing the three environments) using bbmap v38.87 ^14^ and CoverM v0.6.1. Viral MAGs were included in the mapping analysis to enable accurate placement of reads, however, they were not considered in the CoverM analysis since they were extensively analyzed in our previous publication ^13^.

### Gene expression analysis

For each sample, short read poly-A and RiboMinus metatranscriptome data were sequenced and processed by the JGI following SOPs 1027.3, 1082.2, and 1077 (see **Supplemental Information**). Expression quantification was performed using salmon v1.8.0 ^15^ by mapping the processed Poly-A and RiboMinus reads (independently) against the genes predicted from the combined MAG dataset. The Transcripts Per Million (TMP) values produced by Salmon were used for the full community gene expression analysis. To account for the affects that community composition change has on the relative gene expression results when analyzing individual species, the Salmon read counts for genes from *C. merolae* and *G. yellowstonensis* were extracted independently, with their TPM values recalculated using the just these genes. This ensured that when we independently analyzed each species nuclear and organellar gene expression patterns, that the results were free from any community effects.

Orthogroup analysis, conducted using Orthofinder v2.5.4 ^16^, was used to identify the arsenic and mercury detoxification pathway genes in the combined MAG dataset. A parameter sweep was used to identify the optimal inflation value (3.0) for this analysis, using the known Cyanidiophyceae detoxification genes as standards for assessing gene family membership (see **Supplemental Information**). The constructed arsenic and mercury detoxification gene orthogroups had their TPM expression values from the full community expression results analyzed to assess community wide detoxification, enabling the identification of taxonomic groups which pay a disappointedly high role in the expression of particular steps in these pathways.

### Metabolomics analysis

Samples were processed for targeted and untargeted polar metabolomics (positive and negative ionization modes) by the JGI using an integrated extraction and analysis workflow (see **Supplemental Information**). Differentially accumulated metabolites (DAMs) were identified using a two-sided *t*-test, as implemented in the *t.test* function from the stats v4.1.2 R package, with Benjamini & Hochberg adjusted *p*-values computed using the *p.adjust* function (method = “BH”) from the stats v4.1.2 R package.

### Protein structure analysis

Protein structures of ArsM (PDB: 4RSR) and ArsH (PDB: 2q62, just chain A) were downloaded from the Protein Data Bank and visualized in R v4.3.1 using the r3dmol v0.2.0 (https://github.com/swsoyee/r3dmol) package. Key active site residues were identified in the proteins of *G. yellowstonensis* 5572, MtSh, and *G. partita* SAG21.92 homologs and compared among homologs.

## Results

### Analysis of metagenome data from Lemonade Creek

The metagenome read data were mapped against the reference prokaryotic, eukaryotic, and viral MAGs, and the reference nuclear and organelle genomes of *C. merolae* 10D and *G. yellowstonensis* 5587.1 (**Supplemental material**). *C. merolae* was the single most dominant species in all three habitats (**Figs. 1B**), with its plastid genome comprising >50% of the transcripts-per-million (TPM) normalized data in the Creek biofilm and >15% in the Endolithic and Soil habitats (**Figs. 1C**). There was also a significant contribution by *G. yellowstonensis* in the Endolithic and Soil environments, as well as an increase in the proportion of reads derived from other eukaryotic MAGs in the Soil samples (**Supplemental material**). Beyond the algae, a modest contribution was made by bacteria, with <25% of reads being unassigned (i.e., do not map sufficiently well to the assembled reference genomes/MAGs to be classifiable; **Table S1**). The Endolithic and Soil habitats housed a relatively more complex biotic assemblage. More reads derived from Archaea were present in the Endolithic samples, with more bacterial derived reads in both the Endolithic and Soil samples.

The distribution of algae and prokaryotes in Lemonade Creek was studied using sequence co-abundance (**Fig. S1**, **Supplemental material**) and single-nucleotide polymorphism (SNP) data (**Fig. 1D**). These approaches provide consistent results. The SNP analysis showed that both the local *C. merolae* and *G. yellowstonensis* Soil populations have a high degree of distinctness in principal component (PC) 1. Furthermore, whereas the *C. merolae* Creek biofilm and Endolithic populations show a relatively low fixation index (F_ST_) and shared peak regions, the *C. merolae* Soil population has an overall elevated F_ST_ as well as unique peaks, indicating these are distinct populations relative to the other sites (**Figs. S2-S4**), perhaps reflecting species differentiation. Similar trends were observed with the *G. yellowstonensis* populations, although the existence of fewer SNPs in the Creek biofilm population made these calculations less robust (**Figs. S5, S6**).

### Gene expression pattern of YNP MAGs

Poly-A (eukaryote) and RiboMinus (total RNA) metatranscriptomic reads from the Creek biofilm and Soil habitats were mapped against coding sequences (CDS) from the YNP MAGs and the *C. merolae* 10D and *G. yellowstonensis* 5587.1 reference nuclear and organelle genomes to determine relative gene expression levels. Given the observed biotic complexity of the samples and the vastly different abundances of taxa across them, these data are best interpreted as community level shifts in transcription rather than absolute quantification of gene expression in a particular MAG. In the Creek biofilm poly-A transcript data, many reads mapped to bacterial CDS, likely due to contamination [although some bacterial mRNAs are polyadenylated ^17^], which resulted in bacteria comprising a significant fraction (from 4.37% at TP3 to 43.97% at TP1; **Table S2**) of the expressed transcripts (measured as TPM; **Fig. 2A**, left image). As expected, *C. merolae* was the single largest (per genome) contributor to poly-A transcripts (24-98%; **Table S2**), as expected, based on the metagenome data. The Soil poly-A transcript data did not contain a significant number of prokaryotic transcripts (**Fig. 2A**, right image), and *C. merolae* was again dominant in gene expression. Other eukaryotes comprised a significant fraction of the expressed transcripts, with the amoeba *Acanthamoeba* sp. totaling between 3.4-23.4% of the expressed transcripts at each time point (**Fig. 2A** right image; **Table S2**). In the Creek biofilm RiboMinus data, the majority (55.5-96.4%) of transcripts were bacterial derived (**Fig. 2B**, left image; **Table S3**) and the large increase in the relative abundance of *C. merolae* transcripts at TP2 (34.5%) and TP3 (43.7%) compared to TP1 (2.5%) and TP4 (4.2%) was driven completely by plastid genes. In the Soil RiboMinus data, most transcripts are bacterial derived (42.5-74.4%), however, there was an increase in the proportion of archaeal (1.7-5.8%), *G. yellowstonensis* (3-6.6%), and other eukaryote (0.97-2.4%) derived transcriptomes across all time points (**Fig. 2B**, right image; **Table S3**). Still, as for the Creek biofilm data, there is a large contribution to the overall transcript pool by *C. merolae* plastid genes (11.9-41.5%).

**Fig. 2.**
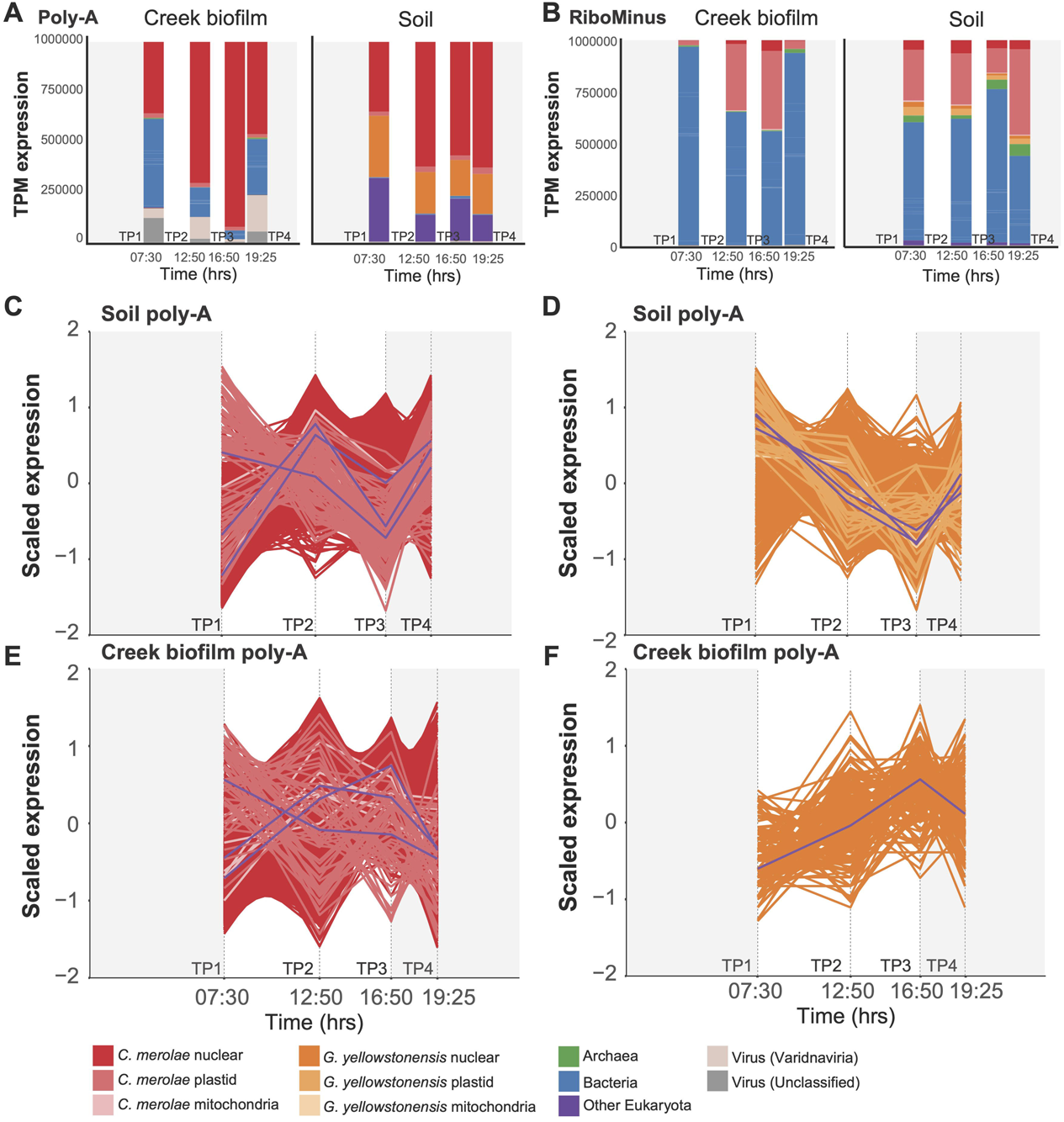
Analysis of YNP metatranscriptome data. Relative abundance (TPM) of (**A**) poly-A and (**B**) RiboMinus metatranscriptome reads mapped against the predicted genes from the assembled YNP metagenome data and reference algal genomes (see legend). Each bar in the graph represents the average value of the n=3 replicates per time point that were analyzed. (**C, D**) Expression patterns of differentially expressed genes in the Soil poly-A libraries. The patterns are presented as log_2_ z-score normalized values. Each point in the line graph represents the average value of the n=4 replicates per time point that were analyzed. The purple lines in the *C. merolae* 10D (**C**) and *G. yellowstonensis* 5587.1 (**D**) panels represent the average expression value of all transcripts at each time point; the three purple lines in each plot represent the average expression of nuclear, plastid, and mitochondrial genes. (**E, F)**. The same analysis of expression patterns done for the Creek biofilm samples.

These data demonstrate that *C. merolae* and many other organisms present in Lemonade Creek follow a diurnal cycle. In the *C. merolae* Soil population, 93.1% of nuclear genes in the poly-A transcript libraries increased significantly (|FC| > 1 and adjusted *p*-value < 0.05) in their relative abundance between TP1 to TP2 (**Fig. 2C**); however, there was a noticeable dip at TP3, before an increase at TP4. *G. yellowstonensis* showed a clear trend of decreasing relative abundance from TP1 to TP3, with an increase at TP4 (**Fig. 2D**). There was a sharper dip at TP3, when *C. merolae* also showed a decrease. This may result from a significant increase in prokaryotic transcription at TP3, or an overall drop in *C. merolae* plastid gene expression coinciding with significantly decreased light levels (**Fig. S7A**), which constitutes a significant proportion of the RiboMinus data (**Fig. 2B**). In the Creek biofilm, 96.2% of *C. merolae* nuclear predicted genes increased significantly in their relative transcript abundance between TP1 and TP2, with none being significantly reduced (**Fig. 2E**). Of the 6,004 *G. yellowstonensis* nuclear genes, 176 (2.9%) significantly increased and 40 (0.7%) significantly decreased (**Fig. 2F**). Because *G. yellowstonensis* comprises a minor proportion of the Creek biofilm metagenome and metatranscriptome samples, the low number of differentially abundant genes in this species is unsurprising.

RiboMinus reads mapped to *C. merolae* genes were extracted and TPM values recalculated to remove the influence that changes in community composition has on the TPM results. These recalculated results show that most expressed *C. merolae* transcripts at each time point are plastid derived (**Fig. S7B, S7C**), with the photosystem II D1 (Q[b]) protein (*psbA*) constituting 39-68% of transcripts in the Creek biofilm (**Fig. S7B**) and 18-45% in the Soil (**Fig. S7C**). In addition, antenna proteins (phycocyanin alpha and beta chain [*cpcA* and *cpcB*] and allophycocyanin alpha and beta chains [*apcA* and *apcB*]) constitute between 7.1-16.2% of total expressed transcripts in the Creek biofilm and 15.3-22.1% in the Soil. In the case of *G. yellowstonensis*, *psbA* constitutes 3.5-19.2% of Creek biofilm and 13.6-21% of Soil transcripts, and antenna proteins constitute 20.2-26.2% of Creek biofilm and 20.7-22.7% of Soil transcripts. There are also several nuclear *G. yellowstonensis* genes with high abundance (>1% total transcripts) in the RiboMinus data, suggesting that this species may be less photosynthetically active than *C. merolae*, relying more on heterotrophy for survival.

### Arsenic and mercury detoxification in Lemonade Creek

Analysis of the diurnal RiboMinus RNA data showed that the relative accumulation pattern of *merA* (OG0000025) in both habitats is dominated by bacteria, with only a minor contribution from *C. merolae* (**Figs. 3C, S8**). The transcript accumulation pattern of *ars* genes is more complex with the *arsC* OG (OG0000808) predominantly expressed by bacteria in the Creek biofilm, and in bacteria and *G. yellowstonensis*, in the Soil. As(III) export potential was more difficult to interpret with these data. Bacterial and *C. merolae arsA* (OG0000997) peaked in their abundance at TP2 and TP3 in the Creek biofilm, whereas *arsB* (OG0000581) was dominated by bacterial taxa with the similar diurnal pattern as *arsA* (these genes are likely to be co-localized in an operon and therefore, co-regulated). The Soil samples were more algae driven, with *G. yellowstonensis* and *C. merolae* OGs dominating the *arsA* and “arsenic transporter” accumulation patterns. *G. yellowstonensis* and bacteria dominating *arsB* (**Fig. S8**). The detected transcripts of *arsM* (OG0000651) in the Creek biofilm derived almost entirely from *C. merolae* (excluding a small bacterial contribution at TP1 and TP4). Furthermore, the change in accumulation of this OG was relatively extreme over the four time points, ranging from < 20 TPM at TP1, to > 200 at TP3, and < 50 TPM at TP4, accounted for by this red alga. The Soil samples showed a similar pattern however, significant contributions were also made by *G. yellowstonensis* and other eukaryotes (**Fig. S8**). Prokaryotes comprised a minor (< 20% total expression) fraction of the detected *arsM* transcripts. Finally, *arsH* (OG0001476) expression in the Creek biofilm was nearly undetectable in the RiboMinus data, with only minor contributions (∼0.02 TPM) from *G. yellowstonensis* at TP3. In Soil, *G. yellowstonensis arsH* was more strongly accumulated, peaking at TP1, and then declined over the day (**Fig. S8**).

**Fig. 3.**
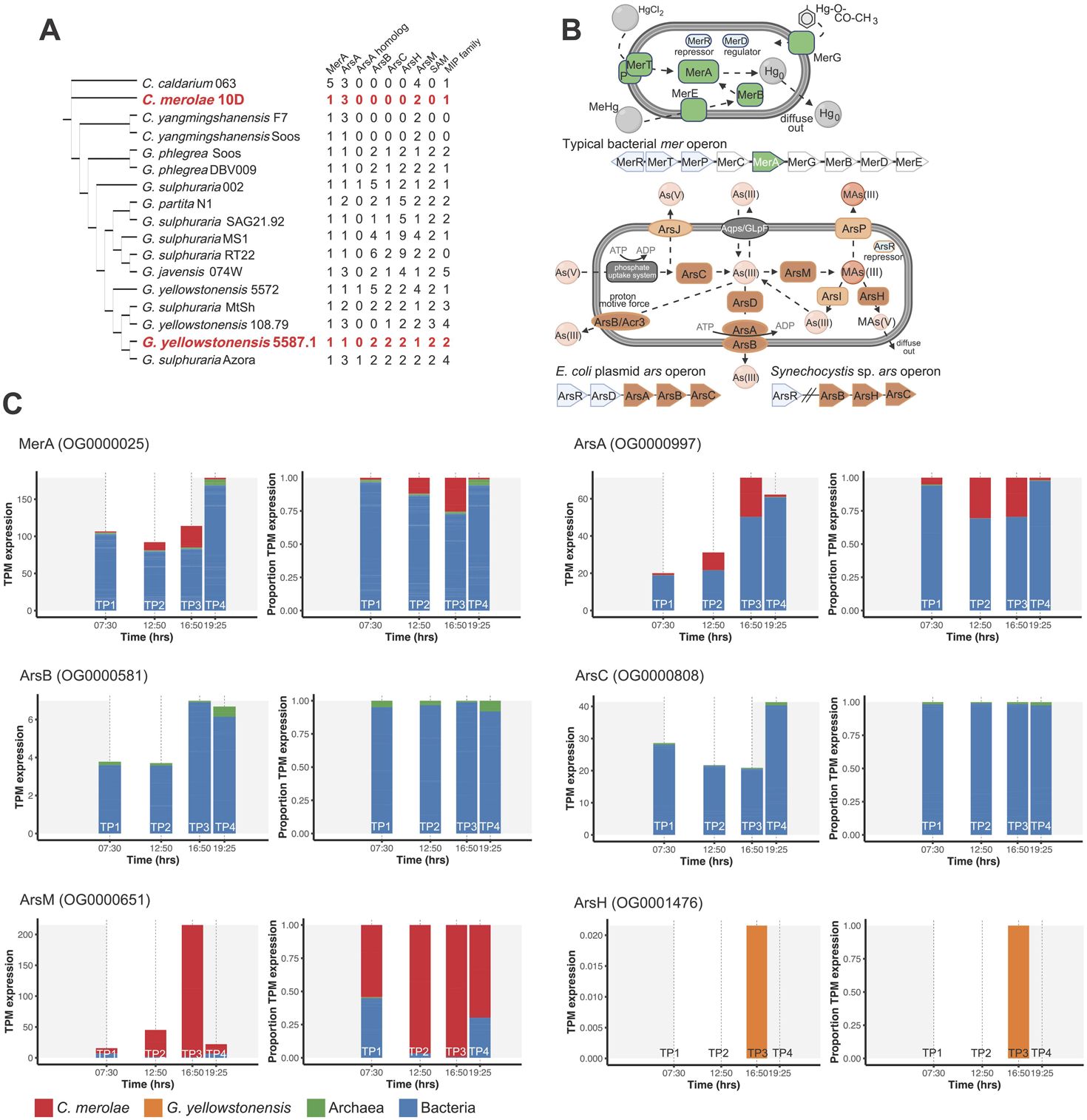
Analysis of *mer* and *ars* genes. **(A)** Distribution and copy number of *mer* and *ars* genes in a canonical phylogeny ^6^ of Cyanidiophyceae. The species (or closely related strains) present in the studied YNP habitats are shown in the red text. **(B)** Schematic prokaryotic cells showing canonical mercury (top) and arsenic (bottom) detoxification pathways. The cells portray the relevant enzymes, however, not all prokaryotes encode all components of these pathways. **(C)** Contribution of taxonomic groups to the arsenic and mercury detoxification pathways in the Creek biofilm samples. The TPM expression values of all genes from an orthogroup identified as containing genes putatively from a specific step in the detoxification pathway is shown as a stacked bar graph (left). The proportion of TPM values contributed by each taxonomic group is shown as a stacked bar graph (right).

### Protein structure

Using existing bacterial protein structures as reference, we analyzed *ars* genes in Cyanidiophyceae to determine whether their functions may have been lost or compromised. For ArsH (beyond *G. partita* SAG21.92), key residues involved in FMN-binding and homotetramerization have been lost (highlighted in **Fig. 4** top). At least one of four ArsM proteins in *G. yellowstonensis* 5572 showed evidence of loss of function; G1662 retains only two of the four conserved cysteines involved in As(III)-binding ^18^ and also lacks most of the residues involved in S-adenosylmethionine (SAM)-binding ^19^ (highlighted in **Fig. 4** bottom).

**Fig. 4.**
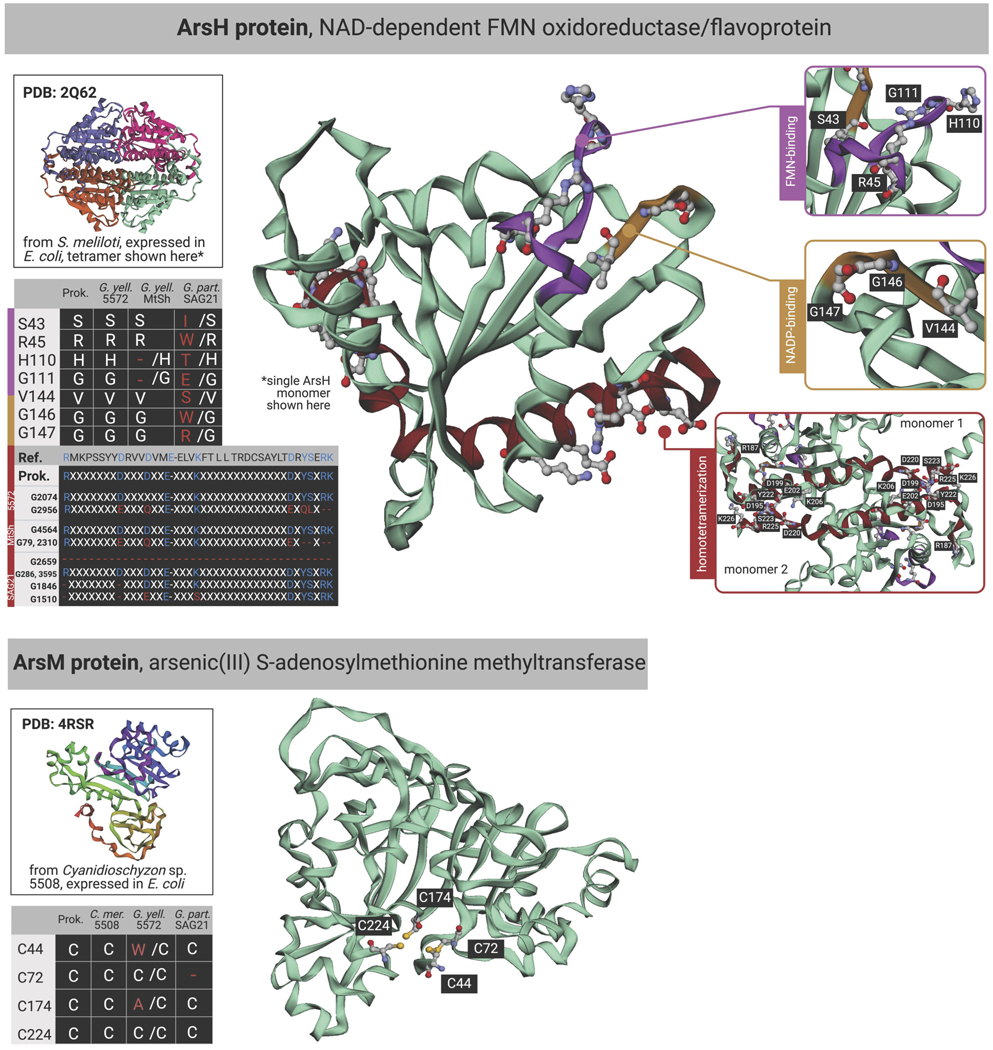
Loss of conserved residues in Cyanidiophyceae ArsH and ArsM. The reference bacterial structures are shown for *arsH* and *arsM*, as are alignments and locations of conserved site changes in the structures that are present in the sequences of the proteins in *G. yellowstonensis* 5572, MtSh, and *G. partita* SAG21.92. These data suggest that the algal *ars* gene original functions may be compromised or lost (**Supplemental material**). Made in Biorender.

### Shifts in metabolism in Lemonade Creek over the diurnal cycle

Untargeted and targeted metabolomic data were generated from the Creek biofilm and Soil samples using pellet (intracellular) and supernatant (extracellular) extracts under positive and negative ionization modes. A total of 26,631 untargeted metabolite features were identified across both modes (19,026, positive mode and 7,605, negative mode). The largest number of differentially accumulated metabolites (DAMs) in the Creek biofilm samples (|FC| > 1, adjusted *p*-value < 0.05) was between TP1 (prior to sunrise) and TP2 (midday) (**Table S4**). There were 3,490 up-regulated and 687 down-regulated features between TP1 and TP2 in the Creek biofilm samples, all other combinations had approximately < 100 DAMs. There were fewer DAMs in the Soil samples due to lack of data from TP1. The untargeted metabolite features (i.e., with putative metabolite identifications) with higher abundance values tended to peak at TP2 or TP3. This pattern with the highest intracellular accumulation at TP1 was observed in the Creek biofilm pellet for a large group of purines and glycosylamines, which likely represent the remnants of cell cycle progression, as indicated by the metatranscriptome data (**Fig. 5A**). However, together with nicotinic acid (precursor of NAD^+^ and NADP^+^) and panthotenic acid (essential cofactor for synthesis of proteins, carbohydrates, and fats) ^20^, they may also indicate increased primary metabolism. Later in the day (TP2 - TP4), an increase in the abundance of osmo-protectants such as trigonelline, marked a change in internal or external solute concentration ^21^. Whereas betaine and choline (see Creek biofilm data) have been described as effective osmo- and thermo-protectants ^22^, carnitine may also play a role as a final electron acceptor in bacteria ^23^.

**Fig. 5.**
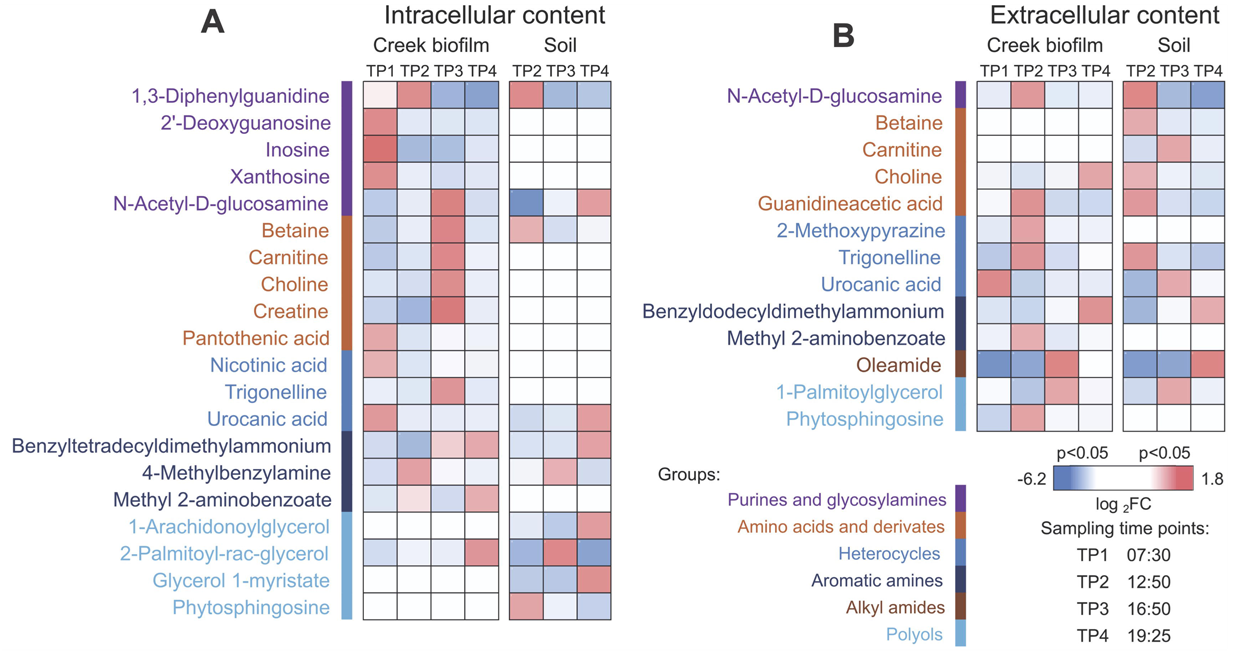
Heatmap of the differentially abundant metabolites in the Creek biofilm and Soil. Pellet collected after the sampling process represents the intracellular content **(A)**, whereas supernatant represents the extracellular content **(B)**. Metabolites were analyzed and annotated *via* the feature-based molecular networking algorithms of the Global Natural Products Social Molecular Networking (GNPS) platform ^42^. Features with a cosine similarity score > 0.72 were utilized for comparisons. Different groups of metabolites are labeled with different colors. The color intensity of each field represents log_2_ FC of the peak area mean in ratio to the overall average for each metabolite in the respective environment. Only significant observations are displayed (ANOVA, *p* < 0.05).

Not surprisingly, a relatively large number of heterocyclic metabolites were found in the extracellular environment of the biofilm, with some that produce foul odors (e.g., 2-methoxypyrazine) (**Fig. 5B**). Urocanic acid (UA) (e.g., Creek biofilm) has previously been described as an algicidal substance secreted by a strain of *Bacillus* sp. ^24^. Interestingly, the points of lowest expression in the *G. yellowstonensis* poly-A RNA data coincided with the highest abundance of UA. Similarly, oleamide, a long-chain amide, has been implicated in the growth inhibition of *Microcystis aeruginosa*, a bloom forming, toxin-producing cyanobacterium ^25^. Oleamides (abundant at TP3 and TP4 in the Creek biofilm and Soil, respectively) have also been described as natural antifouling agents in benthic biofilms that can be synthesized by bacteria or fungi ^26^. In addition, aromatic amines may act as sunscreens, protecting the biofilm from excessive radiation. Benzoates were highly abundant in both Creek biofilm and Soil. The increased abundance of methylated forms suggests they may be produced and secreted by the more abundant bacterial phyla. The highly electrophilic nature of the aromatic ring gives these arenes the ability to act as effective UV screening agents as well as light absorbers within the cell or the biofilm ^27^.

Regarding the targeted metabolomic data, after processing the 109 and 131 metabolites identified under positive and negative ionization modes, respectively, the data were reduced to a unique set of 173 compounds. Abundance changes in this limited number of validated metabolites (using authentic standards) supported insights gained from the untargeted data and metatranscriptomics. For example, in the Creek biofilm intracellular content, the accumulation of sucrose (**Fig. S11**) which is likely to be the major product of photosynthesis in this environment, was opposite of trehalose and the monosaccharides (i.e., sucrose reaches it maximum intensity at TP3, whereas trehalose and the monosaccharides reached their minimum at TP2 and TP3, respectively). This suggests that the sucrose produced by *C. merolae* during the day is being converted to trehalose and (likely) glucose during the night by heterotrophs. These patterns are present, but less pronounced in the Creek biofilm extracellular data (**Fig. S12**).

## Discussion

### *C. merolae* dominates Lemonade Creek biofilms

*C. merolae* is a cell wall-less, obligate photoautotrophic lineage that has a highly reduced genome of size 16.5 Mbp (4,803 nuclear protein coding genes) and is considered to be a specialist, limited to aqueous environments ^28^. Its dominance in the these environments suggests that *C. merolae* outcompetes the physiologically more flexible local *Galdieria* (*G. yellowstonensis*) populations that have the ability to use over 50 sources of carbon for energy ^29^. These data also provide a potential explanation for the larger *Galdieria* gene inventory (e.g., 6,004 nuclear protein-coding genes in *G. yellowstonensis* 5587.1) and their heterotrophic capacity. Namely, these algae retain the ability to use different carbon sources during the dark, which may avoid them competing directly with *C. merolae* which dominates the period of photosynthesis-based energy production in these YNP habitats. SNP analysis demonstrates that despite their close proximity (i.e., within cm), there is significant spatial partitioning of Lemonade Creek biota, which we have previously demonstrated with viruses at this geothermal feature ^13^.

The complexity of the microbial community in these environments and the relative nature of sequencing data, make it challenging to disentangle differential regulation of genes in a particular organism, and the overall shift in the composition of the community. That is, the apparent increase in gene expression in *C. merolae* could be a result of true upregulation of gene expression in this species, the downregulation of genes in other organisms in the community, or a mixture of both processes (the latter being more likely). Regardless of the mechanism, the prokaryote and eukaryote species in these communities clearly follow a diurnal cycle. This result is confirmed by the large contribution of *C. merolae* plastid genes to the RiboMinus transcriptome pool in both environments (**Fig. 2B**). The clear abundance of *C. merolae* plastid genes (particularly the *psbA* and the antenna proteins, which often comprised most of the expressed transcripts) in both the total and species-specific expression results (**Figs. 2B, S7B, S7C**) and lack of other photosynthetic taxa in the sequence data demonstrate that it is the dominant primary producer in both the Creek biofilm and Soil habitats. In contrast, the mixotrophic *G. yellowstonensis* shows relatively greater transcript abundance at night (**Fig. 2D**) and lower plastid (particularly, *psbA*) transcript abundance (**Figs. S7D, S7E**), suggesting that it is more active during the dark phase. This pattern is supported by the untargeted and targeted metabolomic data, which showed that the largest number of DAMs in the Creek biofilm is between TP1 and TP2 (**Table S4**), consistent with strong, light-driven diurnal cycling in these habitats. It also provides insights into the complex biochemical interactions at these sites which supports the long term microbial stability of these environments ^30^.

This pattern of gene expression is consistent with previous lab-based work with synchronized *C. merolae* 10D cultures over the diurnal cycle that showed photosynthetic activity for these obligate phototrophs to peak at midday. Cell cycle progression (G1-S transition) occurred at night to minimize DNA damage caused by redox stress ^29,31^. The latter aspect is critical at the study site where TP2 (midday) light levels (2,003 mmol m^-2^ s^-1^) at Lemonade Creek were one to two orders of magnitude greater than the other times when samples were collected on October 10, 2021 (**Fig. S7A**). However, actual levels in the Creek biofilm and Soil were likely lower due to self-shading. For more details, see **Supplemental material**.

### Arsenic and mercury detoxification pathways are split across multiple taxonomic domains

The geothermal springs in YNP contain significant levels of mercury ^32,33^, particularly in acidic features ^34^, including Lemonade Creek ^35^. Acidic environments favor the occurrence of Hg^2+^ ^33^ suggesting that low pH may have influenced the evolution of *merA* in YNP. A single copy of this detoxification gene is present in both *G. yellowstonensis* and *C. merolae*. In contrast to *merA*, most *ars* genes are present in multiple copies (**Fig. 3A**). MerA encodes reduction of Hg^2+^ to the volatile Hg^0^ that passively leaves the cell ^36^ (**Fig. 3B**), providing resistance. These genes originated from bacteria *via* HGT and share high sequence similarity to prokaryotic MerA, retaining the conserved cysteine residues essential for activity ^37^. Consistent with this idea, MerA in *G. yellowstonensis* 5572 and *G. partita* SAG21.92 (inferred using gene expression analysis) rapidly detoxifies mercury in unialgal culture ^38^. The expression of *merA* in the Creek biofilm and Soil habitats is dominated by bacteria, with only a minor contribution from *C. merolae* (**Figs. 3C, S8**).

Both *G. yellowstonensis* 5587.1 and *C. merolae* 10D encode various *ars* genes, yet the compositions (**Fig. 3A**) and encoded functions differ (see below). And in contrast to *merA* (above), most *ars* genes in these algae are present in multiple copies, all of which were acquired in Cyanidiophyceae *via* HGT (**Fig. 3A**). In the Creek biofilm, based on *arsC* expression levels, As(V) to As(III) conversion by ArsC is a bacterial function, decreasing slightly at TP2 and TP3, and possibly interpreted as shifts in community composition over the diurnal cycle. *C. merolae arsM* expression (encodes for methylation of As(III)) peaks at TP3, which is when this species is most photosynthetically active (inferred using plastid gene expression data; **Fig. 2B**) MAs(III) to MAs(V) conversion, as inferred by *arsH* expression, appears to be absent in this environment. This could be explained by trimethylarsine gas production by ArsM and its spontaneous oxidation to trimethyarsine oxide (TMAsO) ^39^, which is essentially inert, obviating the need for ArsH. The As(III) efflux genes are predominantly expressed by bacteria, with only the ArsA/ATPase family being expressed at TP2 and TP3 by *C. merolae* ^40^. It is worth noting that the *arsC* and *arsM* gene families are broadly distributed across MAGs in our analysis, thus, their expression above detectable levels by specific groups in this environment clearly demonstrates the partitioning of this detoxification pathway across taxonomic domains. The *arsH* gene family is not very broadly distributed, suggesting that it is not an essential part of the arsenic detoxification pathway in the Creek biofilm, apparent in the low relative gene abundance values. The more biotically diverse soil habitat follows a similar trend, however, the Cyanidiophyceae are more dominant contributors to arsenic detoxification, with *C. merolae* again acting as the key methylator of As(III) (**Fig. S8**). Whereas the interaction of *ars* gene expression (and their protein products) across different lineages is inferred here, it is noteworthy that both habitats show internally consistent (albeit different) patterns for this pathway. In the Creek biofilm, all genes except *arsC* show peak relative accumulation at TP3 (early dusk), whereas in Soil, these patterns are more varied with several peaking at TP1 (at or before sunrise). These data suggest a strong functional connection between *ars* gene expression, consistent with a complementary, community-wide detoxification response in both YNP habitats.

Given the extensive primary literature characterizing different *ars* genes, these sequences are ideal for testing the integrated HGT model (IHM) for eukaryotes ^38^. The IHM posits that when conditions that initially favor HGT fixation diminish or disappear (e.g., due to toxin absence or the evolution of community detoxification) foreign gene(s) may survive if they are integrated into a broader stress (or other) responses that favor retention (**Supplemental material**). This hypothesis was built using gene co-expression data for the five *arsH* genes in *G. partita* SAG21.92 which showed them to be strongly linked to clusters of co-expressed genes encoding protein translation and photosynthesis ^38^. The IHM contrasts with the standard model of HGT, as for *mer* genes, whereby this sequence has retained its original function for hundreds of millions of years ^38^. Analysis of *ars* gene protein structure point to a potentially significant outcome of HGT and complex community interactions in eukaryotes: foreign genes may survive *via* duplication, divergence, and putative neofunctionalization that allows them to persist long-term. This is demonstrated by the loss of key functional residues in many ArsH and ArsM gene copies (**Fig. 4**).

## Conclusions

Geothermal habitats in Lemonade Creek, YNP were used to investigate microbial interactions and their impact on gene expression over the diurnal cycle and on longer-term genome evolution. Our work leads to four major insights: 1) the omics data demonstrate that Cyanidiophyceae are the primary drivers of life in these highly acidic geothermal habitats with *C. merolae* as the dominant phototroph (**Figs. 1B, 2, 5**). 2) The cell and diurnal phototrophy-heterotrophy cycles are strongly shaped by local conditions, with shifts in species composition occurring at the scale of centimeters (**Fig. 1D**). 3) Arsenic and mercury detoxification are a “community affair” with members of the three domains of life sharing these duties (**Fig. 3**). *C. merolae* plays a pivotal role as the dominant producer of methylarsenicals that, if they remain as highly toxic trivalent species [e.g., MAs(III)], may regulate local biodiversity *via* microbial warfare across short distances. If, however, they are spontaneously oxidized to inert pentavalent species (e.g., TMAsO), this would play a key role in arsenic detoxification in Lemonade Creek. 4) Community based arsenic detoxification has allowed algal *ars* genes to undergo duplication and sequence divergence, potentially taking on novel functions (**Fig. 4**), as predicted by the IHM. This model is akin to the Black Queen Hypothesis, whereby community interactions (e.g., provision of public goods [metabolites]) can drive prokaryote genome reduction ^41^. If the IHM applies more generally, then highly specialized genes derived *via* HGT and their duplicated copies, may have been the source of novel functions that supported the early evolution of eukaryotes.

## Data availability

All experimental data pertinent to this manuscript is accessible for review, either in the manuscript, in a public database, or as material uploaded with the manuscript as additional files for review purposes. The metagenome and metatranscriptome data associated with this project are available under the Umbrella BioProject link https://www.ncbi.nlm.nih.gov/bioproject/PRJNA1135266, Case CAS-1342124-B7J1V9 - National Library of Medicine Customer Service confirmation TRACKING:000435001319490. The assembled non-redundant MAGs and their associated predicted genes and annotations are available from https://zenodo.org/doi/10.5281/zenodo.12710562. Targeted and untargeted polar metabolomics results are available from https://zenodo.org/doi/10.5281/zenodo.12709950. Feature based molecular networking results for the metabolomics data are available at the Global Natural Products Social Molecular Networking (GNPS) site https://gnps.ucsd.edu/ProteoSAFe/status.jsp?task=8449ead7d6b94e208ceccee688909d83 for the positive mode data and at https://gnps.ucsd.edu/ProteoSAFe/status.jsp?task=013e5b18bc804c9ab5a0f494e297ca73 for the negative mode data. The raw metabolomics data are also available with the MassIVE accession number MSV000095232 and at https://massive.ucsd.edu/ProteoSAFe/dataset.jsp?task=7b95c3e1865444ebbc60d1316517d7f0.

## Acknowledgements

Sampling at Lemonade Creek was conducted and authorized by the NPS research permit YELL-2022-SCI-5364 granted to TRM for conducting research in Yellowstone National Park, USA. JVE was supported by a grant from the National Aeronautics and Space Administration Future Investigators in NASA Earth and Space Science and Technology (80NSSC19K1542). DB and TGS were supported by a grant from the National Aeronautics and Space Administration (80NSSC19K0462). DB was supported by a grant from the National Institute of Food and Agriculture-United States Department of Agriculture (NJ01180). TRM was supported by grants from the National Aeronautics and Space Administration (80NSSC21K0487) and Montana Agricultural Experiment Station (911230). The work (proposal: 10.46936/10.25585/60000481) conducted by the U.S. Department of Energy Joint Genome Institute (https://ror.org/04xm1d337), a DOE Office of Science User Facility, is supported by the Office of Science of the U.S. Department of Energy operated under Contract No. DE-AC02-05CH11231. AO was supported by a grant from the UKRI Biotechnology and Biological Sciences Research Council Norwich Research Park Biosciences Doctoral Training Partnership (BB/T008717/1). TM was supported by a grant from the School of Environmental Sciences, University of East Anglia. HSY was supported by grants from the National Research Foundation of Korea NRF-(2022R1A2B5B03002312, NRF-2022R1A5A1031361). APMW and PW was supported by a grant from the Deutsche Forschungsgemeinschaft (DFG, German Research Foundation) through CRC1535 ‘Microbial Networks’ (Project ID: 458090666). APMW was supported by a grant from the Ministry of Culture and Science of the State of North Rhine-Westphalia, project “Profilbildung 2022 ACCeSS”.

## Author contributions

Conceptualization: DB, TM, JVE, TGS, SLL, HSY

Methodology: TGS, TM, JVE, DB, MY, WC, MC, JG, TRN, BPB, KBL, KB, IVG

Investigation: TGS, JVE, YL, HK, CHC, EC, PW, AO, BPB, KBL, SYM, MY

Visualization: TGS, JVE, TM, CHC, DB

Funding acquisition: DB, JVE, TRN, BPB, KBL, KB, IVG

Project administration: DB, KB, IVG

Supervision: DB, TM, KB, IVG, TM, SYM, MY, APM, HSY

Writing – original draft: TGS, DB

Writing – review & editing: TGS, JVE, TM, DB

---

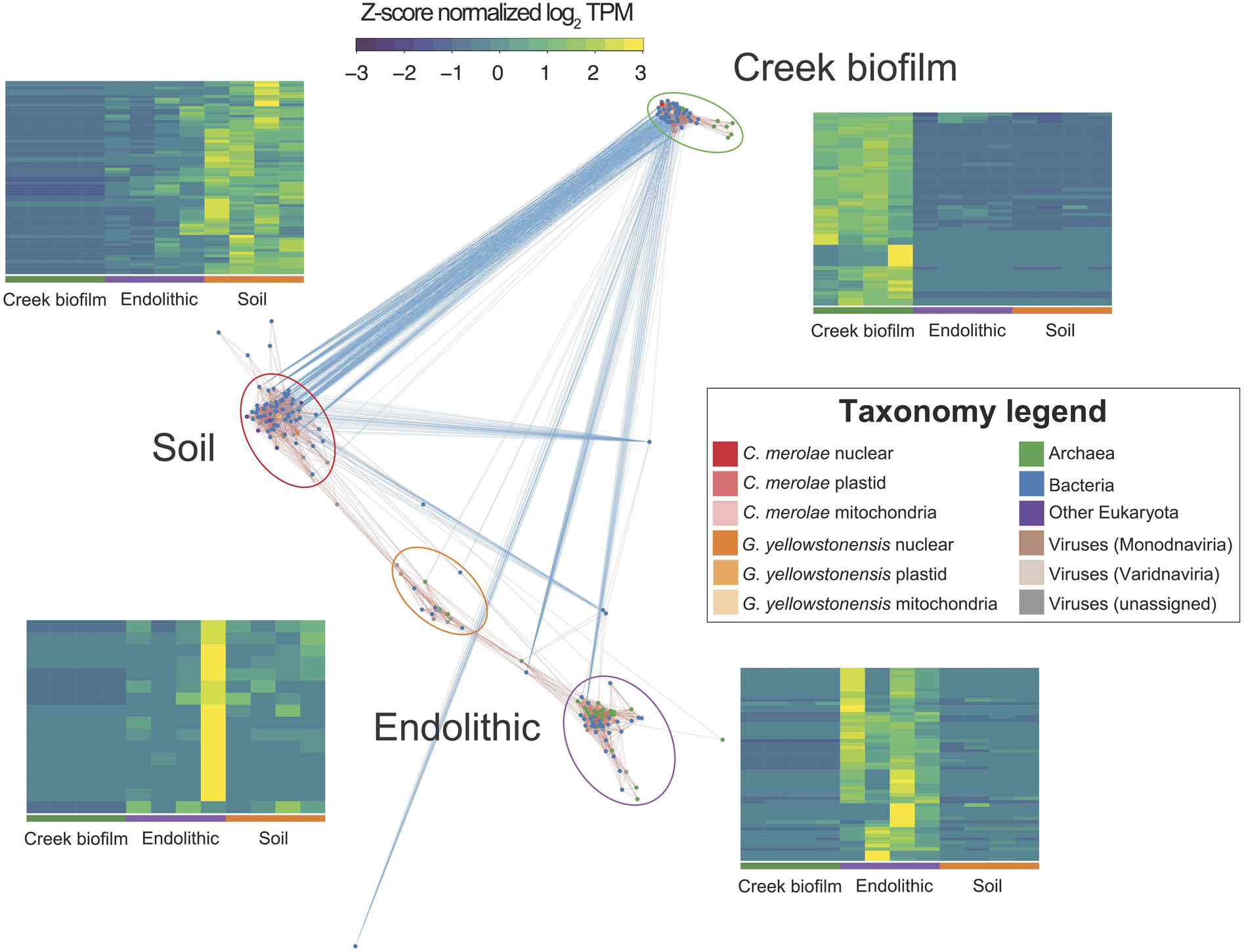

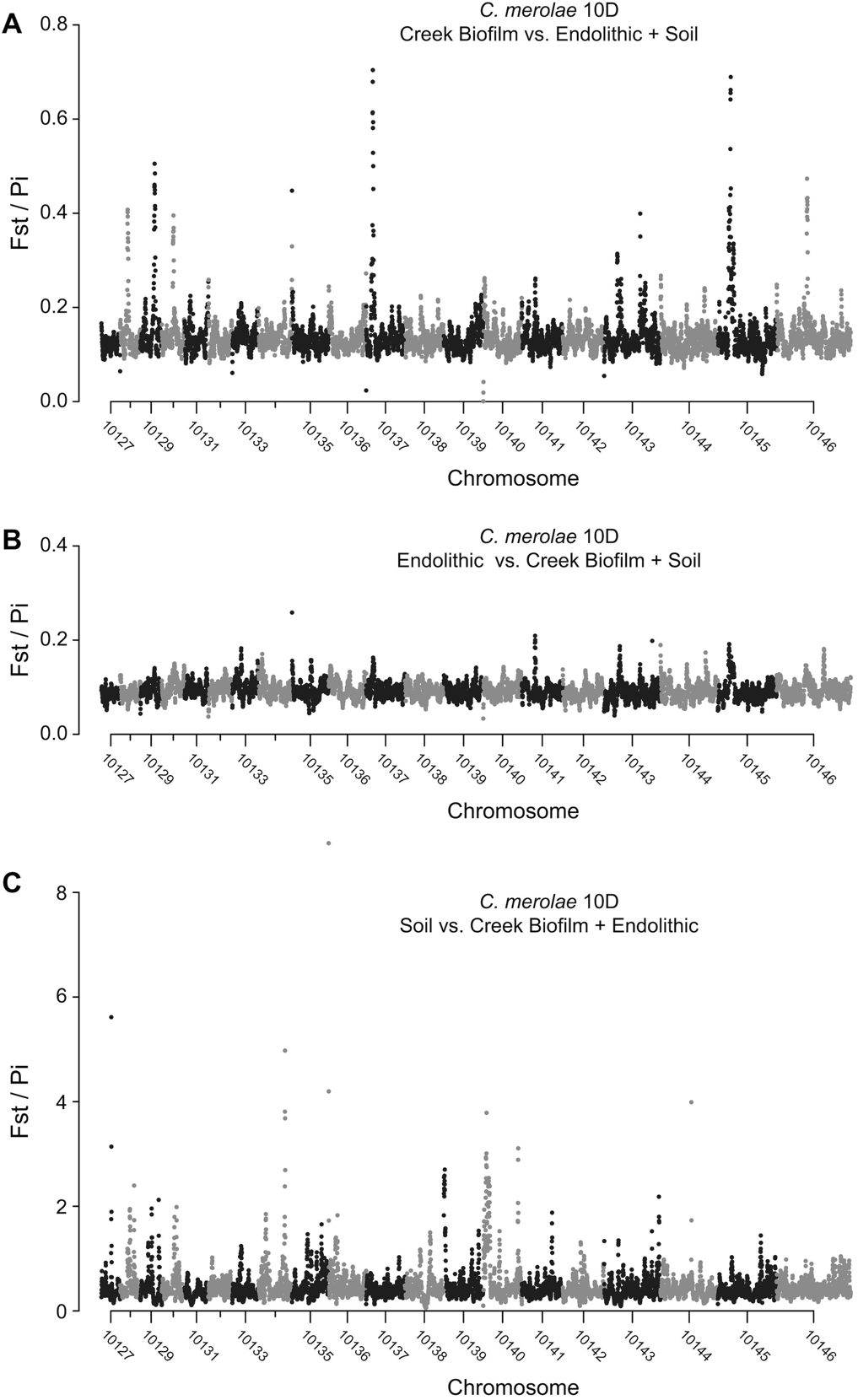

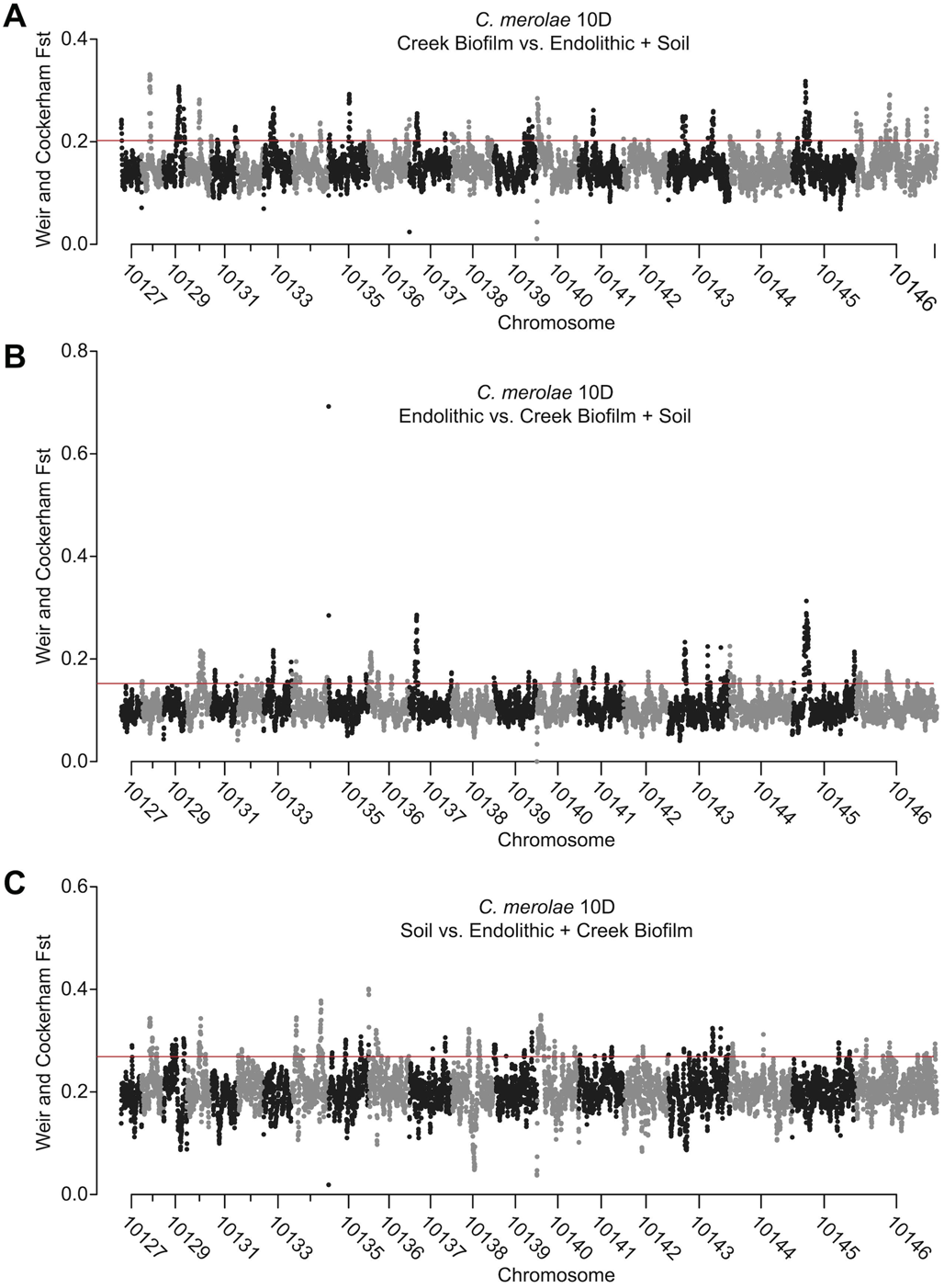

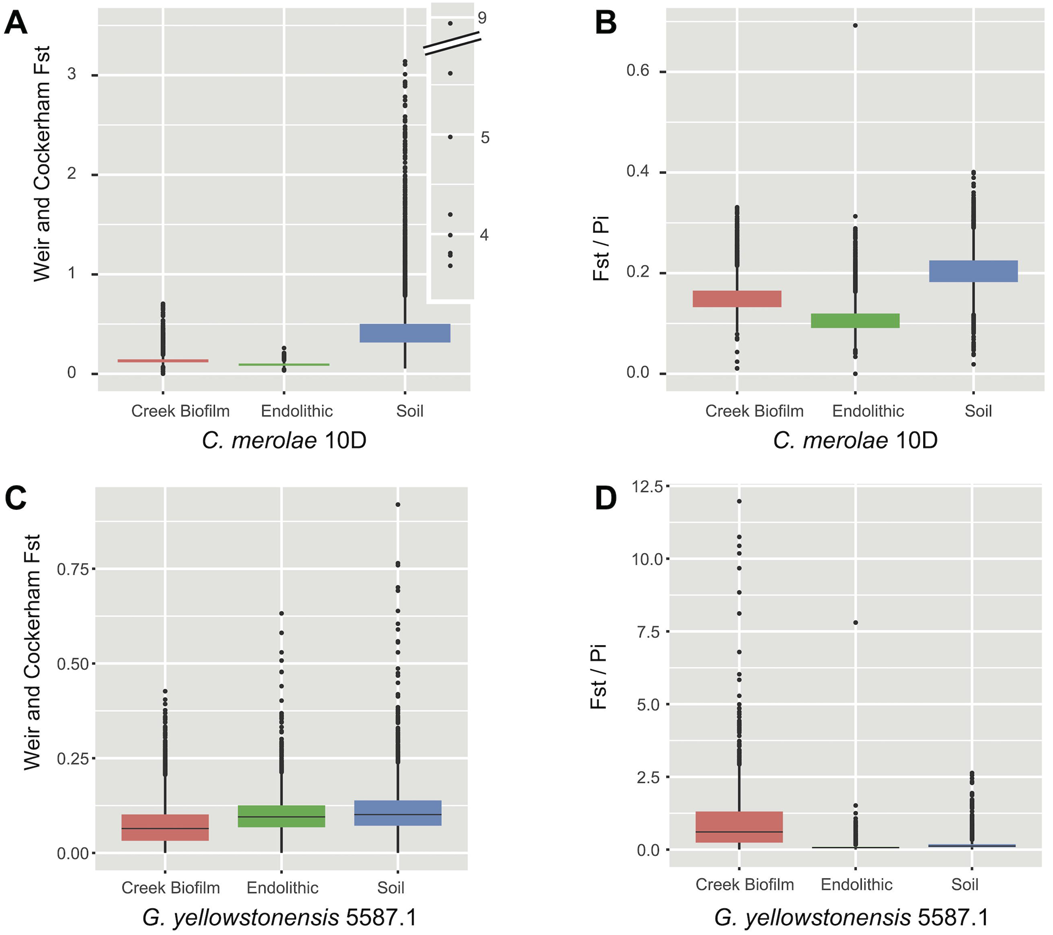

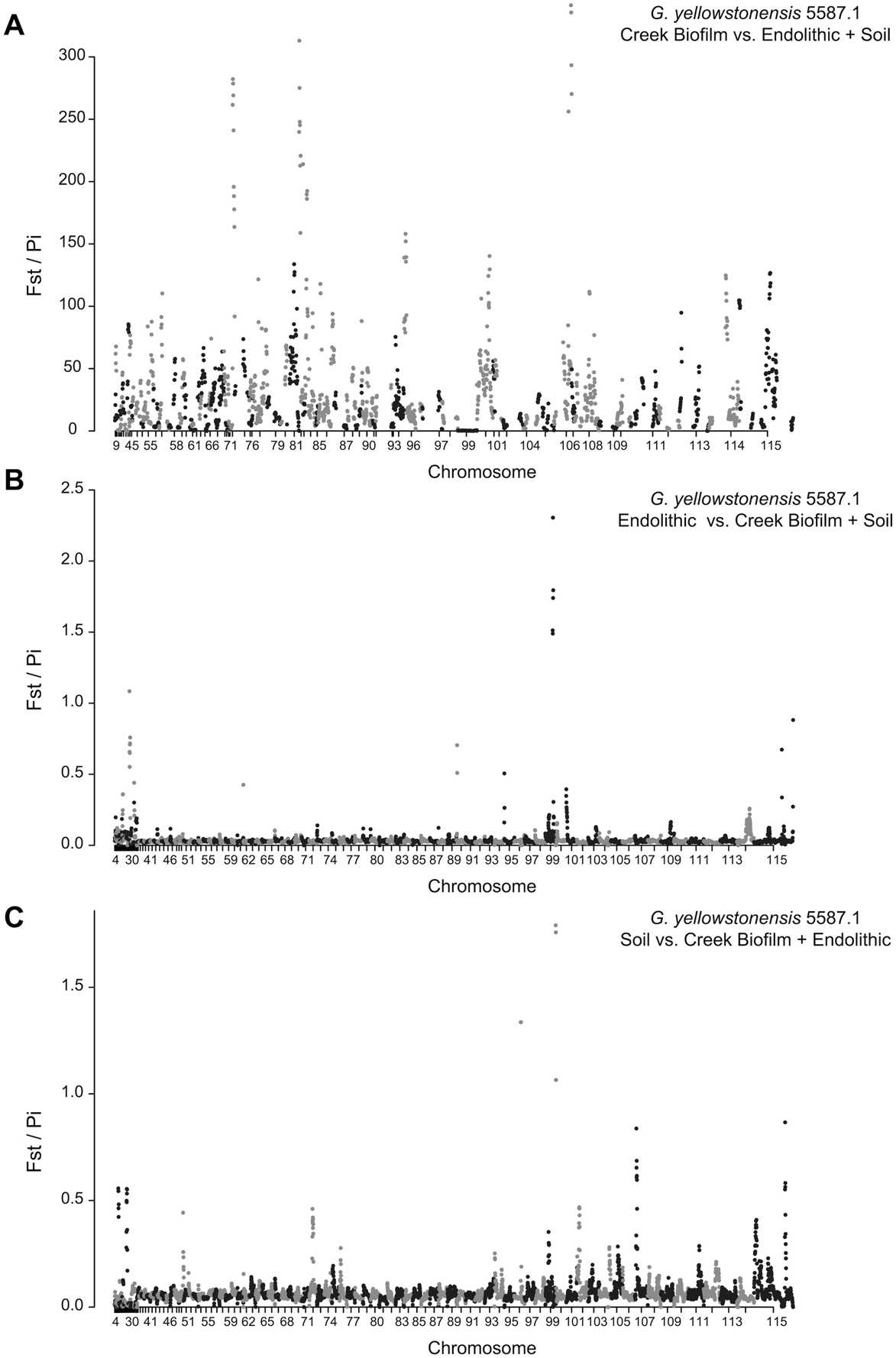

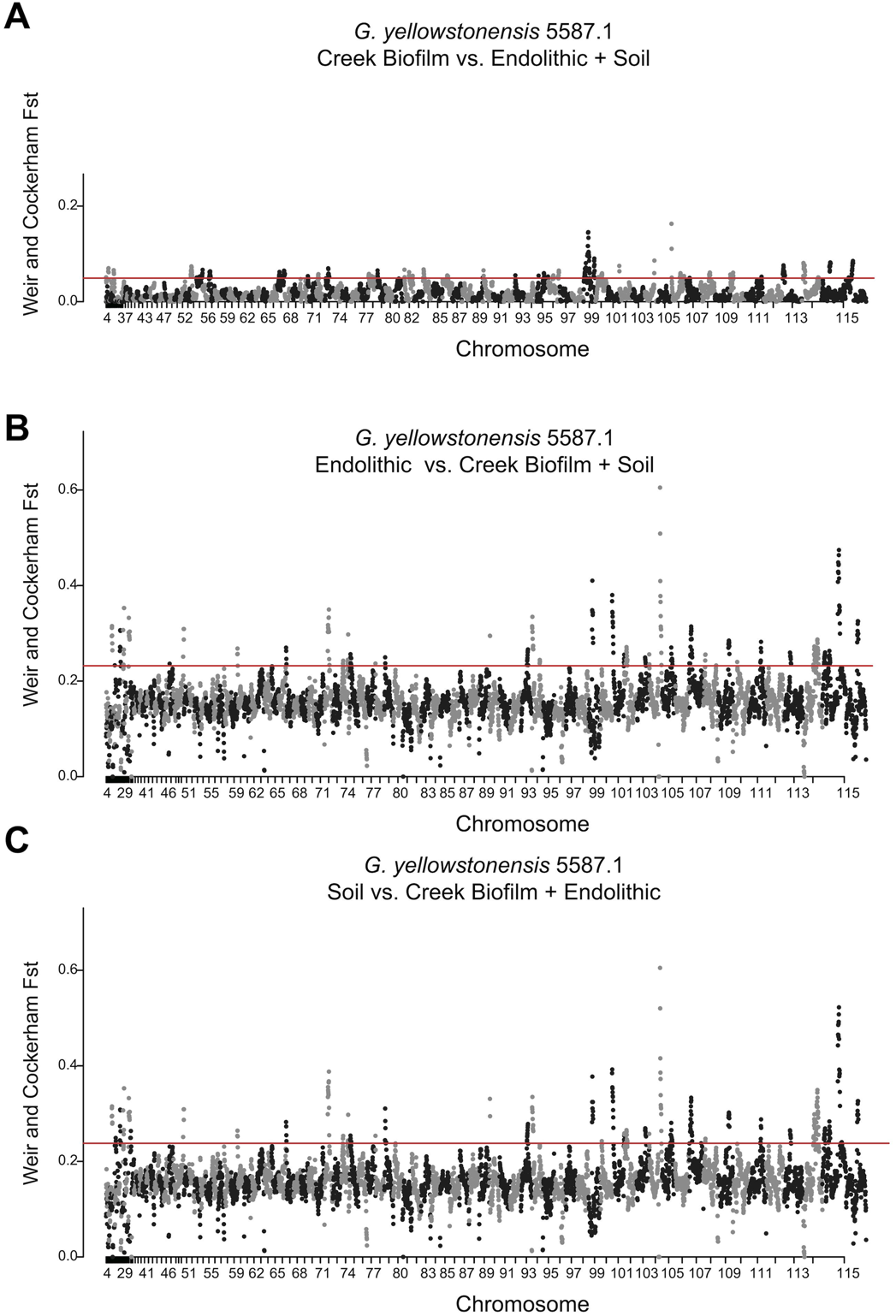

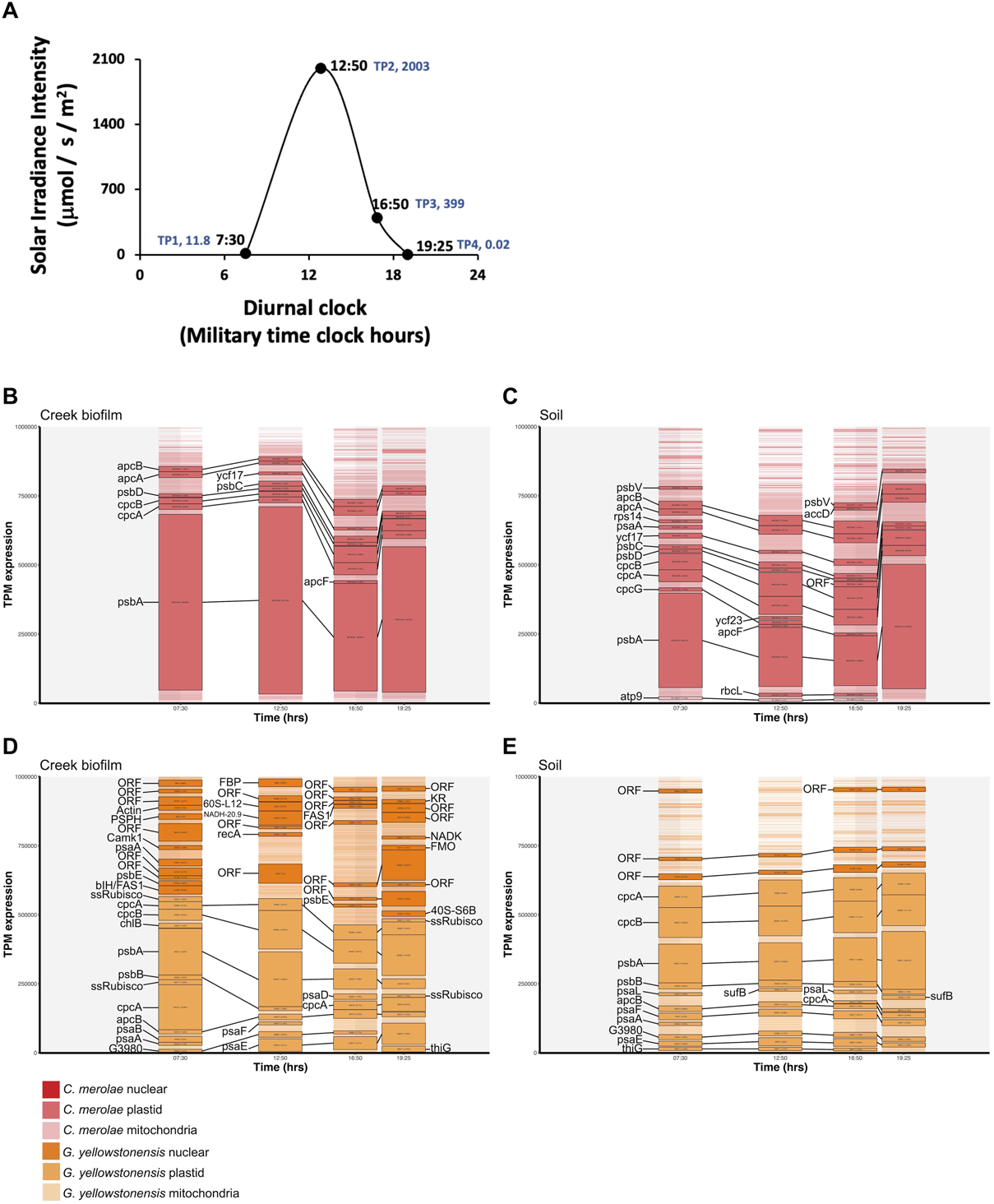

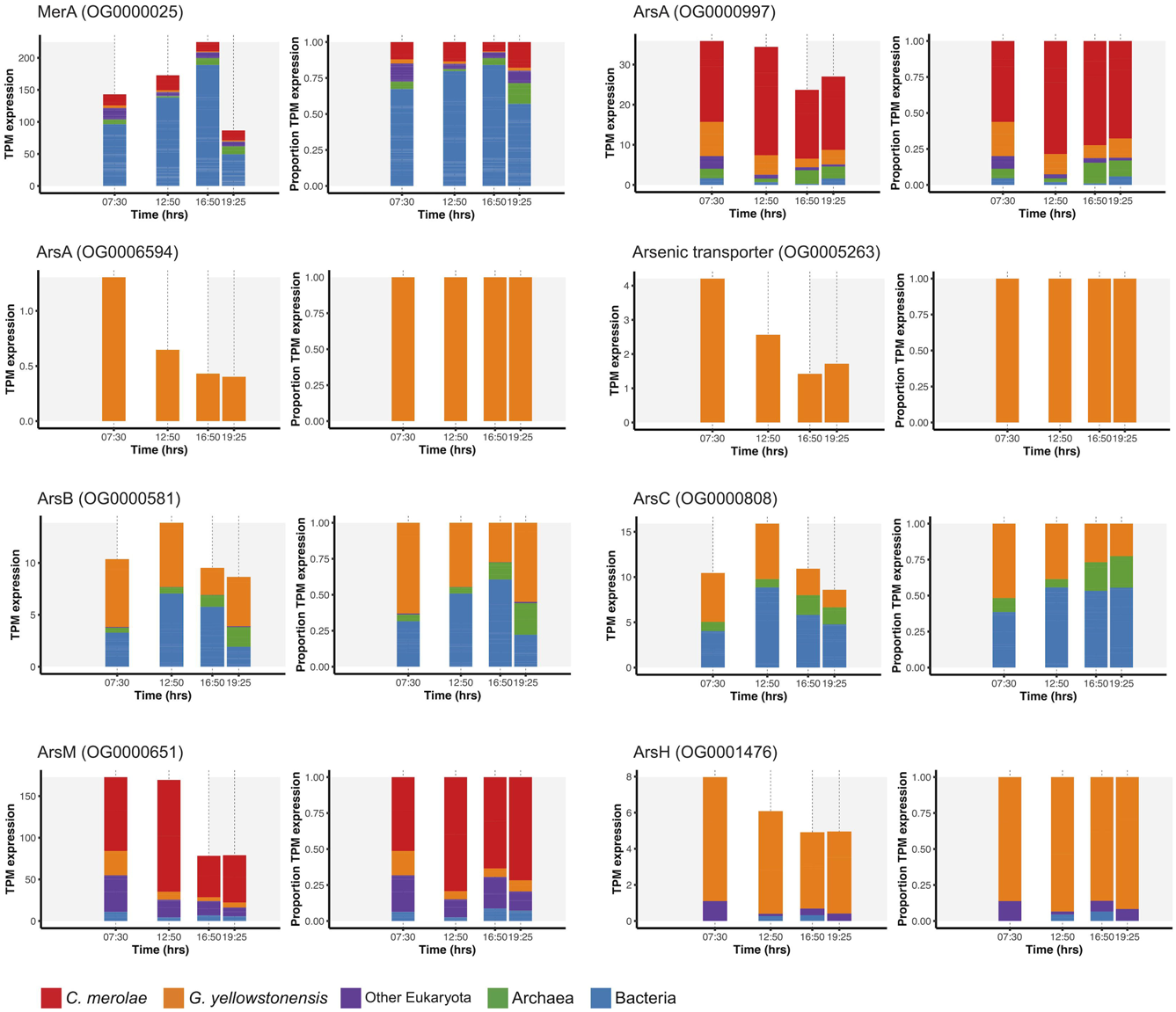

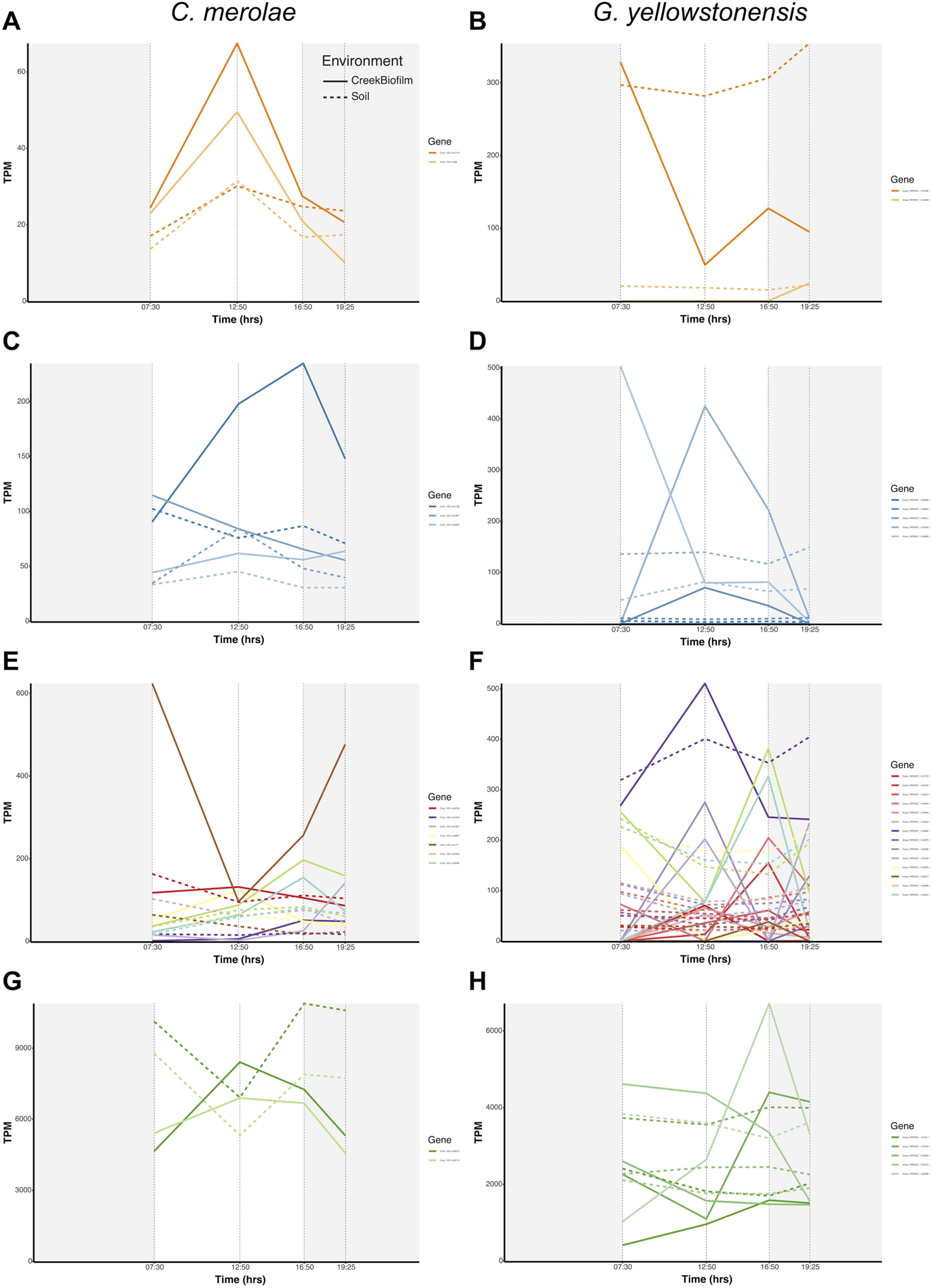

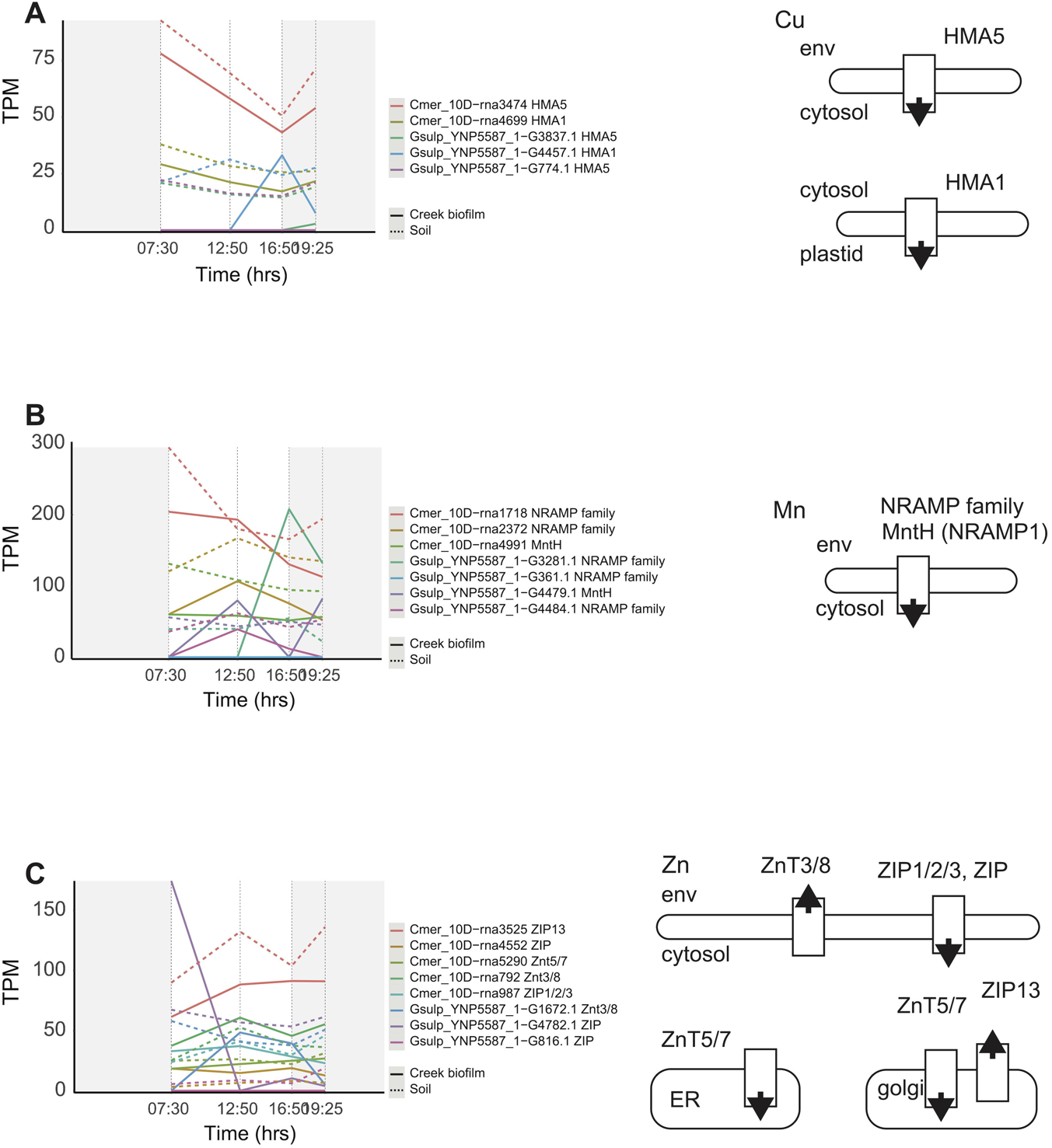

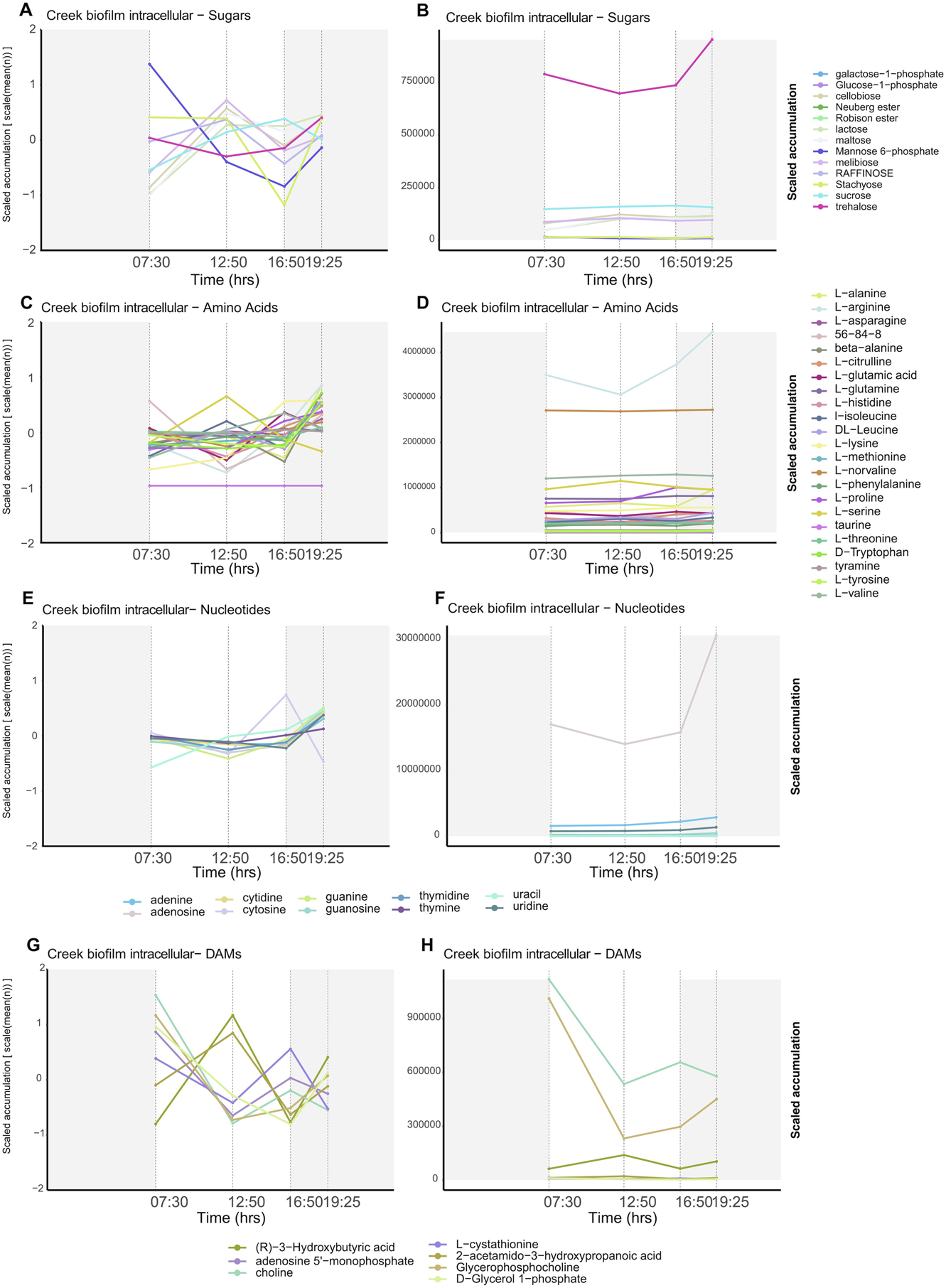

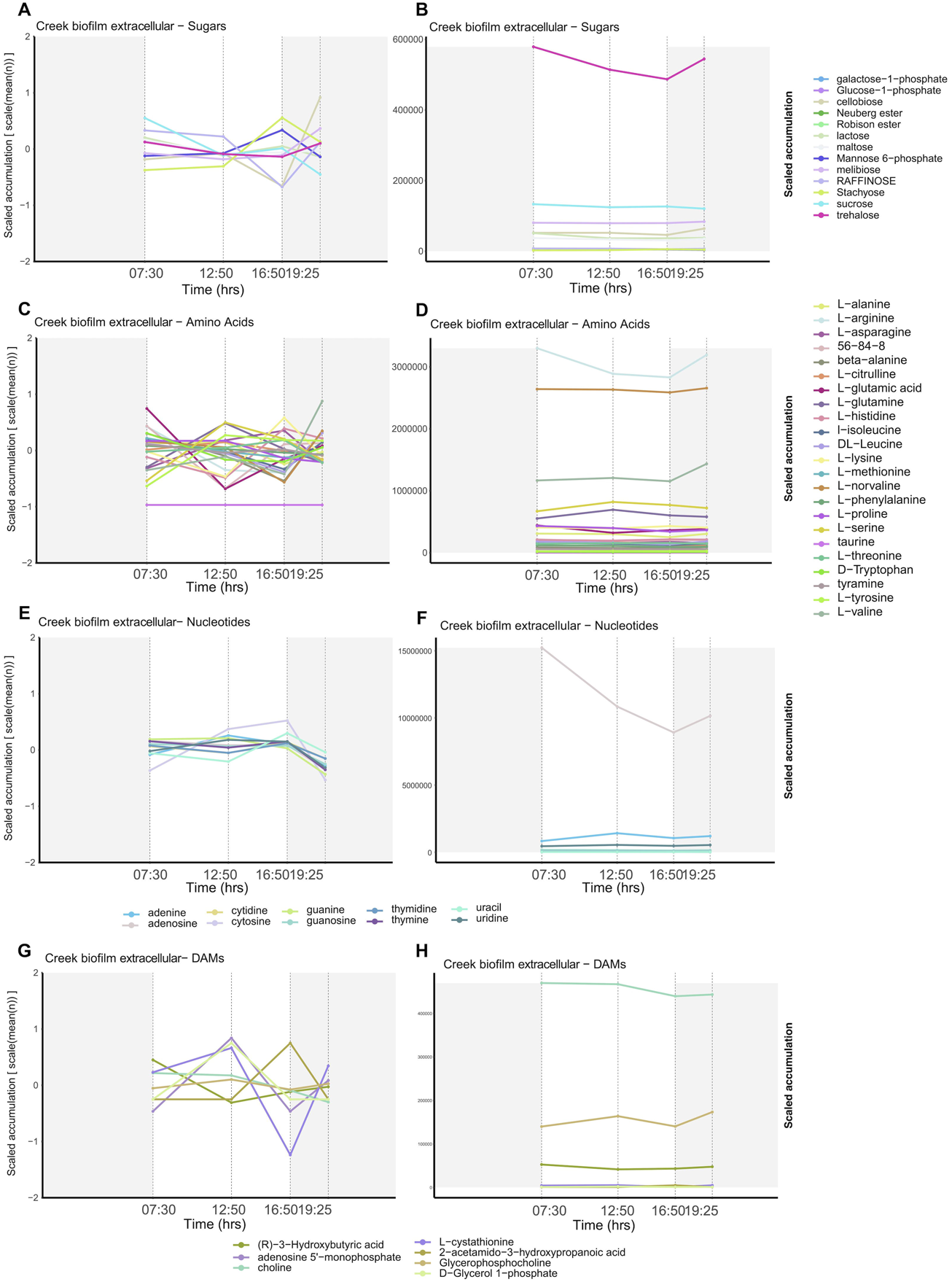

**Supplementary Fig. 13.**
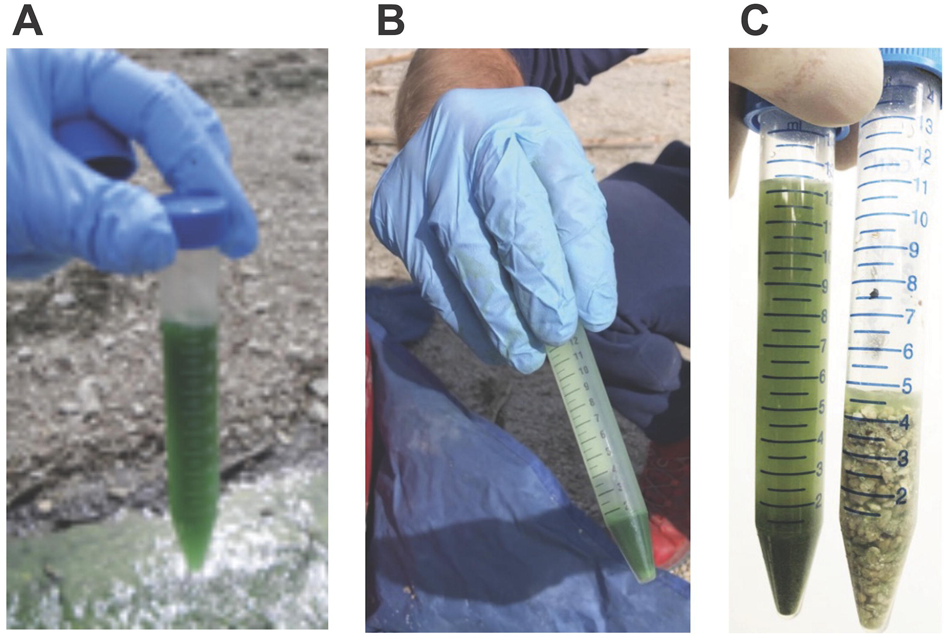
Sampling of YNP habitats at Lemonade Creek. (A) Example of a typical Lemonade Creek bioflim material suspended from rocks and pebbles and decanted to a 15 ml Falcon tube and (B) after centrifugation to pellet biomass and ready for dry-ice ethanol freezing in the field. (C) Preliminary work demonstrating effectiveness of Creek bioflim removal from rocks and pebbles. On the left in the decanted suspension of Creek bioflim biomass stripped from rocks/pebbles by vigorous shaking.

