## Supplemental Information for "Community-wide interactions sustain life in geothermal spring habitats"

**Materials and Methods**

Sampling and field processing

Samples were collected on October 10, 2021, at Lemonade Creek, a geothermal site in Yellowstone National Park (YNP) (44° 48’05” N, 110° 43’44” W). Three distinct environments were sampled at this site, hereafter referred to as “Soil”, “Creek biofilm”, and “Endolithic.”: (1) soils bordering the creek channel, (2) dense lush green biofilms colonizing rocks and pebbles in the creek channel and formed mainly by Cyanidiophyceae, and (3) biomass within rocks sampled within centimeters of the creek flow. During sampling, the creek water was 44°C and pH 2.55, and the soil was 32°C and pH 2.1. These conditions did not change during the diurnal sampling period.

Metagenome sampling

Each environment was sampled at a single time point in quadruplicate (n=4) using gloved hands and sterile tools. For Creek biofilms, manual shaking readily released the algae from the surfaces of the green algae-covered pebbles/rocks. Visual evidence in preliminary work demonstrated that this technique stripped nearly all the green biomass from rocks and pebbles (**Fig. S13**). In the field, rocks and pebbles were transferred to a sterile 50 ml Falcon tube, 5 ml of creek water was added and then manually shaken for ~10 secs. The rocks and pebbles were allowed to settle (~10 secs) and the supernatant was decanted to a 15 ml Falcon tube. In the field, this cell suspension was transferred to a 15 ml Falcon tube and centrifuged at 2100xg for 5 minutes using a benchtop centrifuge powered by a portable generator (both brought to the site). The supernatant was discarded, and the pellet saved (**Fig. S13**). Preliminary testing showed that centrifugation pelleted all algae and was sufficient to pellet bacterial cells with density properties like *Escherichia coli*.

Soil samples were taken by scraping the upper ~0.5 cm of surface soil, carefully avoiding the soil below the discrete algal layer. Endolithic material was acquired by breaking off chips of rock found along the creek edge. Immediately after acquisition, all samples were frozen in an ethanol + dry ice bath, kept on dry ice during transport, and then stored at -80°C.

Metatranscriptome and metabolome sampling

The YNP sampling permit restricted the number of Endolithic samples that could be taken, therefore only a single time point was taken for the Endolithic metagenomic work and this habitat was not included in the metatranscriptomic and metabolomic work. Sampling of Creek biofilms and Soils occurred at four time points (TP1, commencing at sunrise, 07:30; TP2, ~ midday 12:50; TP3, early dusk, 16:50, and TP4, complete darkness, 19:25), with n=4 at each TP. Selection of sampling times assessed the response of Cyanidiophyceae and accompanying Bacteria/Archaea to a range of solar irradiance levels regardless of specific time of day or time interval between samplings.

Creek biofilm metatranscriptomics: The same protocol described above for metagenome sampling was used, except that the biofilm biomass was suspended from the rocks/pebbles using 5 ml of physiologic saline (0.85% NaCl) instead of creek water as a further precaution to avoid cell lysis to preserve cellular nucleic acids. Total elapsed time from stream removal to immersion in dry ice-ethanol bath was ~6-7 min.

Creek biofilm metabolomics: The sampling protocol was like metagenomics except the biofilm material was resuspended from the rocks and pebbles using ultrapure water to prepare the sample for the metabolomics analysis and instrumentation. After centrifugation, the resulting supernatant was transferred to a sterile 15 ml Falcon tube and referred to as “extracellular metabolites”, whereas the cell pellet was referred to as the “intracellular metabolites”. The goal of this step was to separate metabolites within algal and bacterial cells from those assumed to be synthesized and released by the microflora in situ, while recognizing that there may be some cell lysis due to suspending the biofilm cells with ultrapure water (i.e., osmotically sensitive bacteria and archaea). Both types of samples were immediately frozen in in ethanol-dry ice bath and stored on dry ice until transferred to a -80°C freezer.

Soil metatranscriptomics: Sampling was as described for the Soil metagenomic work except that samples (roughly 10 ml by volume) were suspended in 10 ml sterile 0.85% saline. Samples were shaken vigorously for ~ 10 secs., the larger soil mineral material (sand and pebbles) were allowed to settle (~10-15 sec), then the supernatant was decanted to a fresh 15 ml Falcon tube and centrifuged (2100 x g, 5 minutes). The resulting supernatant was discarded, and the pellet (biomass) was flash frozen in an ethanol-dry ice bath and then stored on dry ice until transferred to a -80°C freezer.

Soil metabolomics: Soil samples were aseptically taken by scraping the upper ~0.5 cm of surface soil using sterile sampling tools, transferred to a sterile 15 ml Falcon tube, carefully avoiding the soil below the discrete algal layer. The approximately 10 ml volume of soil was vigorously mixed with 10 ml of ultrapure water (~10 sec), the sand and pebbles were allowed to briefly settle (10-15 sec), and then the suspension was transferred to a 15 ml Falcon tube and centrifuged (2100xg for 5 min). Like the Creek biofilm metabolomics samples, the resulting supernatant (extracellular metabolites) was transferred to a sterile 15 ml Falcon tube, whereas the pelleted material was saved as intracellular metabolites. Both types of samples were flash frozen in ethanol – dry ice bath and then stored on dry ice until transferred to a -80°C freezer.

Lab processing and DNA extraction

Soil and Creek biofilm samples were homogenized *via* bead-beating (vortexed at the maximum speed for 10 min) with 0.5mm silica beads. For Endolithic samples, fragments of rock with visible biomass (e.g., green color) were put inside sterile Ziploc bags and then manually pulverized with a metal mallet hitting the outside of the bag. Once reduced to millimeter-sized grains, they were subjected to the same bead-beating protocol as described above. Total DNA was extracted from the samples with the Qiagen DNeasy PowerSoil Pro kit (Hilden, Germany). Samples were checked for quality, purity, and concentration using the Nanodrop 3000C and Qubit 2.0 and stored at -80°C. These samples were shipped on dry ice to the DOE Joint Genome Institute (JGI) for sequencing.

Metagenome sequencing and assembly

The DNA samples were sequenced by the DOE Joint Genome Institute (JGI). Libraries were created following SOP 1007.3 - LC – using Illumina Nextera with size selection. Tubes containing 0.715 ng of genomic DNA were tagmented using the Nextera XT kit (Illumina) and added JGI Unique dual indices by 9 cycles of PCR. The amplified DNA fragments were size selected with double SPRI using TotalPure NGS beads (Omega Bio-tek). Sequencing followed SOP 1082.2 using the NovaSeq platform. The libraries were quantified using the KAPA Biosystems' next-generation sequencing library qPCR kit and run on a Roche LightCycler 480 real-time PCR instrument. Sequencing of the flowcell was performed on the Illumina NovaSeq sequencer using NovaSeq XP V1.5 reagent kits, S4 flowcell, following a 2x151 indexed run recipe (number of reads and bases sequenced for each sample are listed in **Table S5**).

BBDuk v38.94 ^1^ was used to remove contaminants, trim reads that contained adapter sequences and G homopolymers of size 5 or more at read ends and right quality trim reads where quality drops to 0. BBDuk was used to remove reads that contained 4 or more 'N' bases, had an average quality score across the read less than 3 or had a minimum length <= 51 bp or 33% of the full read length. Reads mapped with BBMap ^1^ to masked human, cat, dog, and mouse references at 93% identity were removed from downstream analysis, as were reads that aligned to common microbial contaminants (following SOP 1077). Reads retained after filtering were corrected using bbcms v38.90 (‘-Xmx100g mincount=2 highcountfraction=0.6’) and assembled independently (i.e., the corrected reads from each library were assembled separately) using spades v3.15.2 (‘-m 2000 --only-assembler -k 33,55,77,99,127 --meta’; **Table S6**).

Alignment of metagenome reads against the assemblies

For each metagenome assembly, the corrected reads from all metagenomes were aligned separately using BBMap v38.87 ^1^ (‘ambiguous=random rgid=filename’) and sorted using samtools sort v1.11 ^2^. The jgi_summarize_bam_contig_depths tool from MetaBAT2 v2.15 ^3^ was used to calculate contig read mapping depth, using the separate bam files produced from mapping of the corrected reads as input. Read mapping rates for each pair-wise combination of metagenomes is listed in **Table S7**.

Identification of sample specific Cyanidiophyceae metagenome contigs

A total of 16 Cyanidiophyceae genomes ^4–6^ (**Table S8**) were acquired from public databases and were compared against the assembled metagenome from each sample to assist with identification of contigs originating from the red algal species in each sample. The assembled metagenomes were compared against a database consisting of the 16 Cyanidiophyceae genomes using BLASTn v 2.10.1 (‘-max_target_seqs 2000 -evalue *E* ≤ 10^-10^ -dust no -soft_masking false’); repeat masking in BLAST was disabled to allow for better alignment of repetitive regions between the two datasets. Only hits which had >90% identity were retained for downstream analysis. Hits that overlapped along each scaffold were merged into a single feature and the coverage of each scaffold by the resulting merged BLASTn hit features was calculated using bedtools v2.29.2 (i.e., using the sort, merge, and coverage commands). Scaffolds with >90% of their bases covered by merged BLASTn hits were classified as high confidence Cyanidiophyceae sequences. The putative origin of the high confidence Cyanidiophyceae scaffolds was assigned using the top hit (i.e., the hit with the highest bitscore) for each scaffold. That is, a scaffold was assumed to have originated from the same (or a close relative) species as its top hit. If a scaffold had multiple hits with equally high bitscores then it was assumed to have originated from multiple species; because this occurred extremely rarely and the potential ambiguity was not considered a problem.

The assembly quality and completeness of the extracted high confidence Cyanidiophyceae metagenome scaffolds and Cyanidiophyceae reference genomes was assessed using the statswrapper.sh script from BBMap v38.87 ^1^ and BUSCO v5.0.0 ^7^ (‘--mode genome’) using the ‘eukaryota_odb10’ dataset (**Table S9**). The Average Nucleotide Identity (ANI) between the extracted Cyanidiophyceae metagenome scaffolds and Cyanidiophyceae reference genomes was calculated by PYANI v0.2.11 ^8^ using the ‘ANIm’ method (**Table S10**).

Construction of sample-specific prokaryotic MAGs

First, the high confidence Cyanidiophyceae sequences (those with >90% coverage by BLASTn hits with >90% identity) were removed from each of the metagenome assemblies to produce a set of putative non-Cyanidiophyceae scaffolds that were used as the basis for prokaryotic binning. Contig depth information, that was previously calculated using the jgi_summarize_bam_contig_depths tool, was extracted for the selected non-Cyanidiophyceae scaffolds. The non-Cyanidiophyceae scaffolds were initially binned using three tools: (1) GroopM2 v2.0.0 ^9^ (commands: ‘parse --singlem’, ‘core -s 200000’, and ‘extract’ [default parameters]; using SingleM v0.13.2 [https://github.com/wwood/singlem]), (2) MaxBin2 v2.2.7 ^10^ (‘-markerset 40 -min_contig_length 1000’), and MetaBAT2 v2.15 ^3^ (‘--seed 42424242 --minContig 2500 --minClsSize 200000’). For each metagenome, the sequences and contig depths of the non-Cyanidiophyceae scaffolds were used for MaxBin2 and MetaBAT2 binning; the sequences of the non-Cyanidiophyceae scaffolds and unfiltered BAM files (produced by alignment of the corrected reads against the full metagenome assemblies using BBMap) were used for GroopM2 binning. The three sets of bins (one from each tool) were combined, and a non-redundant set produced (per metagenome sample), using the MetaWRAP v1.3.2 ‘bin_refinement’ command ^11^ (‘-c 50 -x 10’). The quality and completeness of the GroopM2, MaxBin2, MetaBAT2, and MetaWRAP bins was assessed using the statswrapper.sh script from BBMap v38.87 and CheckM v1.2.1 (‘lineage_wf --tab_table’; using checkm_data_2015_01_16).

Each of the bins produced by MetaWRAP were filtered for putative contaminant scaffolds (i.e., scaffolds from different prokaryote lineages) using MAGpurify v2.1.2 ^12^, MDMcleaner v0.8.3 ^13^, and whokaryote v0.0.1 ^14^ (**Table S11**). MAGpurify was run using the ‘phylo-markers’, ‘clade-markers’, ‘known-contam’, ‘tetra-freq --weighted-mean’, and ‘gc-content --weighted-mean’ commands on the selected bins, with any scaffolds that were identified as putative contaminants by any of the previous commands removed using the ‘clean-bin’ command. MDMcleaner was run using the “clean” command (default parameters) on the cleaned bins from MAGpurify. Finally, whokaryote (‘--minsize 500’) was used to remove putative eukaryotic sequences from the cleaned bins from MDMcleaner. As whokaryote has decreased accuracy on scaffolds below 5 Kbp (which made up a significant fraction of some bins), the classification results of MDMcleaner were used to prevent false positives from being erroneously removed; only scaffolds identified by whokaryote as being of eukaryotic origin and had been assigned a 'toplevel_taxlevel' classification of either "None" or "root" by MDMcleaner were removed.

Per-environment co-assembly of non-sample-specific contigs

Reads from contigs in each metagenome sample that were not classified as either high confidence Cyanidiophyceae (using BLASTN [**Table S9**]; described previously) or prokaryotic (from the bins produced by MetaWRAP [**Table S11**], described previously) were extracted (using sambamba v0.8.2, samtools v1.11, and a modified version of the ‘filter_reads_for_bin_reassembly.py’ script from MetaWRAP) and reassembled (using spades v3.15.5; ‘--meta --only-assembler’), combining each of the samples from the same environment together to produce one co-assembly per environment (**Table S12**). The reads used for co-assembly were mapped independently (per sample) against each of the co-assemblies using BBMap v38.87 (‘ambiguous=random rgid=filename’) and sorted using samtools sort v1.11. The jgi_summarize_bam_contig_depths tool from MetaBAT2 v2.15 was used to calculate contig read mapping depth, using the separate bam files produced by BBMap. Any contigs in each of the co-assemblies with >90% coverage by hits with >90% to sequences in the *C. merolae* 10D, *G. yellowstonensis* (formally *G. sulphuraria*) YNP5587_1 and Azora genome assemblies were removed from downstream analysis. This was performed to remove any red algal data that was only recovered in the co-assemblies and not assemblies of the individual metagenome samples. For each co-assembly, the remaining contigs were grouped into bins by MetaBAT2 v2.15 (‘--seed 42424242 --minContig 1500 --minClsSize 1500’) using the associated read mapping depth information for each contig.

Taxonomic classification of contigs from per-environment co-assemblies

Open reading frames (ORFs) were predicted in the contigs of each per-environment co-assembly using prodigal v2.6.3 ^15^ (‘-p meta -f gff’) and compared against the nr database (2022_07) using diamond v2.0.15 ^16^ (‘blastp --iterate --ultra-sensitive’). The top hit returned for each ORF (if one was identified) was used to assign the putative taxonomic provenance of that sequence, with the full lineage information for each top hit retrieved using TaxonKit v0.12.0 ^17^. ORFs without top hits or unknown lineage information we assigned “NA” values. At a given taxonomic rank, the number of ORFs from a given taxon were counted, and the taxon with the most assigned ORFs was used as the putative taxonomic classification of each contig (i.e., a majority vote classification system). If there were equal numbers of ORFs assigned to multiple taxa at a given rank, then the one with the highest average bitscore across the assigned ORFs was selected as the putative taxonomic classification. This process was also performed at different taxonomic ranks and using all ORFs predicted in each bin (i.e., majority classification of a bin using ORFs predicted across all contigs in that bin).

Bins that were classified at the Domain level as “Archaea”, “Bacteria”, or “Eukaryota”, and that had total assembly sizes of over 500 Kbp, 100 Kbp, or 2.5 Mbp (respectively), were extracted for further analysis. The minimum assembly size thresholds were chosen as they are roughly below the sizes of the smallest free-living genomes described for those groups; while it is possible that species with smaller genomes exist for each of these groups, they will likely have highly reduced genomes with few marker genes, making it difficult to assess their quality and to integrate them with the other assembled MAGs. The extracted archaeal and bacterial bins were processed to remove putative contaminant scaffolds and assess completeness using the same workflow described previously.

The completeness of the extracted eukaryotic bins was assessed by BUSCO v5.4.3 ^7^ (‘--mode genome’) using the ‘eukaryota_odb10’ lineage. Bins which had low BUSCO completeness (close to 0%) also had the majority of their ORFs assigned to a range of different taxa at the Phylum level (i.e., the bin appeared to be from a range of organisms), thus these bins were considered low quality and not considered for further analysis. For each of the remaining bins, the lowest ranked taxon name with >50% of the ORFs (identified in that bin) assigned to it was used as the putative taxonomic classification. Additional sequences were recruited to each of the classified bins by combining them (separately) with other bins and unbinned scaffolds in the same per-environment co-assembly which have the same taxonomic classification. Putative contaminant sequences were removed from each of the combined bins using two approaches: (1) scaffolds without the same taxonomic classification as the parent bin, or that were not classified as eukaryotic, were removed; (2) scaffolds predicted by whokaryote v0.0.1 (‘--minsize 500’) as being prokaryotic and that weren’t classified as the same taxa as the parent bin were removed. The combined and filtered bins produced from this workflow were used as the final eukaryotic MAGs in downstream analysis. The assembly statistics of each final eukaryotic MAGs was computed by statswrapper.sh script from BBMap v38.87, the completeness of each MAG was assessed by BUSCO v5.4.3 (‘--mode genome’) using the most appropriate lineage based on the putative taxonomic classification (**Table S13**).

Construction of non-redundant prokaryotic MAGs

The sample-specific prokaryotic MAGs (**Table S11**) and the prokaryotic MAGs derived from the per-environment co-assemblies (described in previous section) were combined into a non-redundant set of Lemonade Creek prokaryotic MAGs using dRep v3.2.2 ^18^ (‘dereplicate --completeness 10 --contamination 25 --P_ani 0.7 --S_ani 0.98’) (**Table S14**). An average nucleotide identity (ANI) of 98% was used for dereplication because these MAGs were to be used for abundance analysis with the available metagenome and metatranscriptome samples and, as suggested by the dREP authors, 98% represents the threshold at which genomes are distinct when mapping short reads. That is, it is challenging to differentiate genomes with above 98% ANI using short-read data, therefore, there would be little benefit when including them in our downstream abundance analysis as we would not be able to reliably differentiate them.

Assembly quality and taxonomic assessment of prokaryotic MAGs

For each prokaryotic MAG (per-sample and non-redundant), assembly statistics were computed using the statswrapper.sh script from BBMap v38.87, completeness and contamination was assessed using CheckM v1.2.1 ^19^ (‘lineage_wf --tab_table’; using database version 2015_01_16), and putative taxonomic provenance was assessed using GTDB-Tk v2.1.1 ^20^ (‘classify_wf’ command; using database version r207_v2) (**Tables S11, S14**). Following the standards set forth by The Genome Standards Consortium ^21^, a MAG was considered to have “High” completeness if it had >90% completeness and <5% contamination, “Medium” completeness if it had ≥50% completeness and <10% contamination, and “Low” completeness if it had <50% completeness and <10% contamination.

Prokaryotic MAG gene prediction and functional annotation

Genes were predicted in the archaeal MAGs using prokka v1.14.6 ^22^ (‘--kingdom Archaea --metagenome --addgenes --addmrna’) and in the bacterial MAGs using bakta v1.5.1 ^23^ (‘--keep-contig-headers’; database version 4.0). Functions were assigned to predicted proteins using eggnog-mapper v2.1.6 ^24,25^ (‘--pfam_realign denovo --report_orthologs --no_file_comments --dbmem’; database version 5.0.2) and InterProScan v5.53-87.0 ^26^ (‘-dp --goterms’) (**Tables S11, S14**).

Eukaryotic MAG gene prediction and functional annotation

Genes were predicted independently for each of the five eukaryotic MAGs constructed from the per-environment co-assemblies (**Table S13**) using the following workflow. Repeats were identified in each eukaryotic MAG using RepeatModeler v2.0.1 ^27^ (‘-LTRStruct’). The RepeatModeler *de novo* repeats were combined with the repeats from Dfam v3.3 ^28^ and the repeats packaged with RepeatMasker to produce a custom repeat family library for each MAG. The MAGs were masked by RepeatMasker v4.1.2-p1 ^29^ (‘-no_is -a -x -gff’) using the custom repeat family library. Given that the eukaryotic MAGs were derived from the Soil co-assembly, the Soil corrected Poly-A selected transcriptome reads were aligned against the MAGs using STAR v2.7.8a ^30^ (‘--limitBAMsortRAM 350000000000 --twopassMode Basic --outSAMtype BAM SortedByCoordinate’). Genes were predicted in each of the MAGs by BRAKER v2.1.6 ^31–40^ (‘--softmasking --gff3 --min_contig=2000’) using the softmasked assemblies produced by RepeatMasker and the aligned transcriptome reads produced by STAR. The only exception to this workflow was the *Stylonychia* sp. MAG which had to have its gene model manually trained (instead of using BRAKER self-training like the other eukaryotic MAGs) due to the unusual genome structure of ciliates combined with their use of an alternative codon translation table ^41,42^ (table=6) that the BRAKER self-training code can't accommodate. Manual training involved extraction of expressed transcripts from the STAR aligned reads using StringTie2 v2.0.6 ^43^ (‘--rf’) to produce genes models and gffread v0.11.6 ^44^ (‘-w’) to extract the transcript sequences. A database of 568,529 ciliate protein sequences were retrieved from NCBI using the search term ‘txid5878[Organism:exp]’ on the 25^th^ Jan 2023. TransDecoder (retrieved from GitHub [<https://github.com/TransDecoder/TransDecoder>] on 26^th^ Jan 2023) was used to identify ORFs in the expressed and extracted transcripts. The ‘TransDecoder.LongOrfs’ script was run using the ‘-G Ciliate’ parameter and ‘TransDecoder.Predict’ script was run using the ‘--single_best_only -G Ciliate’ parameters, with extrinsic homology evidence also provided to the latter script by a DIAMOND v2.0.15 (‘--max-target-seqs 1 --ultra-sensitive --iterate --outfmt 6 --evalue 1e-5’) search against the Ciliate protein database downloaded from NCBI. Next, the existing AUGUSTUS gene model parameters for the ciliate *tetrahymena* was retrained following the protocol by Hoff et al. ^33^ using the gene models of the expressed transcripts with TransDecoder predicted ORFs. Briefly, the ORFs identified by TransDecoder were filtered to remove those without start and stop codons (i.e., were not annotated by TransDecoder as “complete”). Additionally, the remaining proteins were clustered using DIAMOND (performed using the ‘aa2nonred.pl’ script available with BRAKER) to remove sequences with >80% similarity. At each stage of the training process, the ‘--translation_table=6’ parameter was added to the commands specified by Hoff et al. ^33^ to enforce usage of the ciliate codon translation table. Upon completion of the training workflow, BRAKER was run on the *Stylonychia* sp. MAG using the manually trained model, the softmasked genome, and the STAR aligned RNA-seq reads (‘--softmasking --gff3 --min_contig=2000 --translation_table=6 --skipAllTraining --useexisting --extrinsicCfgFiles=BRAKER/Augustus/config/extrinsic/extrinsic.M.RM.E.W.cfg’). If BRAKER predicted multiple isoforms per gene, then only the longest isoform was retained for downstream analysis. The completeness of the predicted genes in MAG was assessed by BUSCO v5.4.3 (‘--mode protein’) using the most appropriate lineage based on the putative taxonomic classification (**Table S13**). Functions were assigned to predicted proteins using eggnog-mapper v2.1.6 (‘--pfam_realign denovo --report_orthologs --no_file_comments --dbmem’; database version 5.0.2) and InterProScan v5.53-87.0 (‘-dp --goterms’).

Metatranscriptome sequencing and assembly

RNA-seq libraries were prepared and sequenced by the DOE Joint Genome Institute (JGI). Library creation was done following SOP 1027.3 Illumina Low Input RNA-seq w/rRNA Depletion (Fast select). The rRNA was removed from 100 ng of total RNA using Qiagen FastSelect 5S/16S/23S for bacterial rRNA depletion (and additional FastSelect plant and/or yeast rRNA depletion) (Qiagen) with RNA blocking oligo technology. The fragmented and rRNA-depleted RNA was reverse transcribed to create first strand cDNA using Illumina TruSeq Stranded mRNA Library prep kit (Illumina) followed by second strand cDNA synthesis which incorporates dUTP to quench the second strand during amplification. The double stranded cDNA fragments were A-tailed and ligated to JGI dual indexed Y-adapters, followed with an enrichment of the library by 10 cycles of PCR. Sequencing was done following SOP 1082.2 using the Illumina NovaSeq platform. The libraries were quantified using the KAPA Biosystems' next-generation sequencing library qPCR kit and run on a Roche LightCycler 480 real-time PCR instrument. Sequencing of the flowcell was performed using NovaSeq XP V1.5 reagent kits, S4 flowcell, following a 2x151 indexed run recipe (number of reads and bases sequenced for each sample are listed in **Table S15**).

BBDuk ^1^ was used to remove contaminants, trim reads that contained adapter sequence and G homopolymers of size 5 or more at the ends of the reads and right quality trim reads where quality drops to 0. BBDuk was used to remove reads that contained 1 or more 'N' bases, had an average quality score across the read less than 10 or had a minimum length <= 51 bp or 33% of the full read length. Reads mapped with BBMap ^1^ to masked human, cat, dog and mouse references at 93% identity were removed from downstream analysis, as were reads that aligned to common microbial contaminants (following SOP 1077). Reads that were identified as known spike-ins and ribosomal RNA (only for the PolyA library) were removed.

Metagenome read mapping and abundance analysis

The abundance of the 174 non-redundant prokaryotic MAGs (**Table S14**), the *C. merolae* 10D nuclear (GCF_000091205.1), plastid (AB002583.1), and mitochondrial (NC_000887.3) reference genomes, *G. yellowstonensis* YNP5587_1 nuclear (4), plastid (GsulpYNP55871_99), and mitochondrial (GsulpYNP55871_27) reference genomes, the five eukaryotic MAGs (**Table S13**), and the 3,679 viral vOTUs ^45^ was quantified across the Soil, Endolithic, and Creek biofilm environments using the 12 sequenced metagenome samples (**Table S5**). The corrected reads from each metagenome sample were aligned against a database consisting of the previously listed reference genomes and MAGs using bbmap v38.87 ^1^ (‘ambiguous=random rgid=filename’), with the resulting aligned reads sorted by coordinates using samtools sort v1.11 ^2^. The coverage of each MAG or reference genome in the combined dataset was quantified using CoverM v0.6.1 (<https://github.com/wwood/CoverM/>; ‘genome --min-read-percent-identity 98 --min-covered-fraction 0 --min-read-aligned-percent 70 --output-format sparse --methods relative_abundance mean trimmed_mean covered_bases variance length count reads_per_base rpkm tpm’), with the 98% minimum read percent identity cutoff chosen as it represents the threshold at which genomes are distinct when mapping short reads, the 0 minimum covered fraction cutoff chosen as we wanted genome/MAGs with no or very low coverage to still be reported in the output (by default genomes with < 10% coverage are excluded from the output), and the 70 percent minimum read aligned percent was to exclude reads with very partial local alignments from affecting the abundance statistic.

Metatranscriptome read mapping and abundance analysis

The abundance of the genes predicted in the 174 non-redundant prokaryotic MAGs (**Table S14**), the *C. merolae* 10D nuclear (GCF_000091205.1), plastid (AB002583.1), and mitochondrial (NC_000887.3) reference genomes, *G. yellowstonensis* YNP5587_1 nuclear ^4^, plastid (GsulpYNP55871_99), and mitochondrial (GsulpYNP55871_27) reference genomes, the five eukaryotic MAGs (**Table S13**), and the 3,679 viral vOTUs ^45^ were quantified across the four time points, in the Soil and Creek biofilm environments, using the RNA-seq data generated by Poly-A selection and RiboMinus library preparation protocols. The coding sequences (CDS) from the genes predicted in the previously mentioned reference set were combined into a single sequence database, which was indexed by salmon v1.8.0 ^46^ (default parameters) using the corresponding reference genome assemblies as decoys. For each of the sequenced transcriptome samples (from each time point, environment, and library preparation method), salmon v1.8.0 (‘quant --validateMappings --seqBias --gcBias --posBias --libType A’) was used for sequence quantification and calculation of the Transcripts Per Million (TMP) statistic (**Table S16**). The expression results from samples from the same environment and library preparation method were combined into a single matrix using salmon before downstream analysis. Each expression matrix (i.e., all time point replicates sequences from an environment produced using a specific library preparation method) was filtered to remove genes which didn’t have ≥ 2 samples with a Counts Per Million (CPM) ≥ 0.5 (i.e., genes with near zero expression across all the samples being considered). The filtered read count matrix was used for differential expression analysis by DESeq2 v1.42.0 ^47^, with each time point compared to all subsequent time points (i.e., time point 1 vs. 2, 3, 4, time point 2 vs. 3, 4, and time point 3 vs. 4). Only genes with an adjusted *p*-value < 0.05 and an absolute log_2_ fold-change > 1 were considered significantly differentially expressed (**Table S17**).

Orthogroup construction and analysis

Orthogroup analysis was used to identify the arsenic and mercury detoxification pathway genes in the non-redundant prokaryote, eukaryote, and viral MAGs. The predicted proteins from the 174 non-redundant prokaryotic MAGs (**Table S14**), the five eukaryotic MAGs (**Table S13**), and the 3,679 viral vOTUs ^45^ were clustered into orthogroups along with proteins from the available Cyanidiophyceae reference genomes (**Table S8**) by Orthofinder v2.5.4 ^48^ (‘-S diamond_ultra_sens’) using the inflation values of 1.5 (default), 2.0, 3.0, 5.0, and 10.0 (**Table S18**). A set of known arsenic and mercury detoxification genes were extracted from the functional annotation of the Cmer_10D, Cmer_SOOS, Gphleg_SOOS, Gsulp_002, Gsulp_074W, Gsulp_5572, Gsulp_Azora, Gsulp_MS1, Gsulp_MtSh, Gsulp_RT22, Gsulp_SAG21, Gsulp_YNP5587_1 reference genomes (**Table S8**) using the phrases “arsen” and “mercu”. Orthogroups constructed using each of the examined inflation values were extracted and manually examined if they contained any of the annotated Cyanidiophyceae arsenic or mercury detoxication genes that we previously extracted. The eggNOG-mapper annotations (described previously) for each of the proteins in the extracted orthogroups were used to assess the specificity of the orthogroups, that is, the uniformity of the eggnog annotations was used to assess if the orthogroup contained proteins with a specific function or if the proteins were a mix of multiple related functions (**Table S19**). An inflation value of 3.0 was chosen as being the optimal parameter for this analysis as gene orthogroups appeared to be relatively uniform in their annotations and higher inflation values did not appear to significantly (if at all) improve the results. Therefore, the orthogroups identified by OrthoFinder using an inflation value of 3.0 were used for downstream analysis.

Metabolite extraction and processing

To extract metabolites for LC-MS, biofilm pellets were first transferred to 2 mL tubes using 1 mL MilliQ water, then all samples (Biofilm pellets - 0.5 to 3 g; Soil pellets – 0.5 – 1.5 g; Biofilm supernatant ~8 mL; Soil supernatant ~8 mL; Creek water, 10 mL) were frozen and lyophilized dry (FreeZone 2.5 Plus, Labconco). To each sample, 100% methanol was added (1 mL for supernatants, Creek water and Soil, 2 mL for Biofilm pellets), vortexed briefly, then sonicated 10 minutes in a water bath. Samples were centrifuged for 5 min at 5000 rpm to pellet debris, then solvent transferred to another tube and dried in a SpeedVac (SPD111V, Thermo Scientific). In preparation for LC-MS analysis, extracts were resuspended in 180 mL methanol containing isotopically labeled internal standards (<https://doi.org/10.1093/plphys/kiae001>) then centrifuge-filtered (0.22 um hydrophilic PVDF membrane, UFC40GV0S, Millipore) prior to transferring to glass LC-MS vials. Due to the low pH of Creek water as well as Soil/Biofilm supernatants (pH 2-3), an additional 3 mL of LC-MS water was added to each sample to test pH and then 0.1 M NaOH added incrementally to adjust sample pH to between 5-7. The pH-adjusted samples were frozen and lyophilized dry, resuspended in 176 mL of 100% methanol before being transferred to glass LC-MS vials.

LC-MS analyses were performed on metabolite extracts using HILICZ chromatography on an Agilent 1290 UHPLC coupled to a Thermo QExactive HF Orbitrap (Thermo Scientific, San Jose, CA) mass spectrometer, using the same methods and instrument settings (<https://doi.org/10.1093/plphys/kiae001>). Sample groups consisted of 4 biological replicates and corresponding extraction controls, with randomization of sample injection order and a blank of 100% methanol between each sample, as well as internal standard and QC mix injections interspersed throughout the run.

To identify metabolites, both targeted and untargeted analysis was performed on the datasets. For targeted analysis, identifications were made based on comparing retention time (RT, exact mass and MS2 fragmentation spectra to that of a database of standards run in-house using the same LC-MS methods. A score ranging from 0 to 3 was given to each feature (unique mz/RT combination) to evaluate the level of confidence in identification. For positive identification, or Level 1, a metabolite had detected m/z < 5ppm (or </-0.001 Da) from theoretical and RT within 0.5 minutes from theoretical (RT-adjusted spectra) compared to the metabolite standard. The highest level of positive identification, Exceeds Level 1 (score of 3), additionally had matching fragmentation spectra when compared to in-house spectra. Other identifications are putative or invalidated by mis-matching fragmentation spectra.

The polar targeted metabolite accumulation results processed such that (1) blank values (i.e., samples where a metabolite was not detected) were filled using a value that was 2/3 the minimum intensity value produced across the whole dataset (each ionization mode was processed separately), and (2) metabolites from the positive and negative ionization modes were combined, keeping, in cases were a metabolites was detected in both ionization modes or with multiple different adducts in the same mode, the metabolite with the highest median intensity across the treatment and control samples. The polar untargeted metabolite accumulation results were processed such that blank values were filled using the same process as the targeted metabolites. The positive and negative ionization modes were combined and since none of the m/z and retention times matched between the two modes, no duplicates were removed. Differentially accumulated metabolites were identified in the processed polar targeted and untargeted datasets using a two-sided *t*-test, as implemented in the *t.test* function from the stats v4.1.2 R package, with Benjamini & Hochberg adjusted *p*-values computed using the *p.adjust* function (method = “BH”) from the stats v4.1.2 R package. VIP scores were calculated using the *mixOmics* v6.18.1 R package ^49^. Targeted metabolites were considered significant if they had an absolute fold change (FC) > 1 and a *p*-value < 0.05, untargeted metabolites were considered significant if they had an absolute fold change (FC) > 1 and an adjusted *p*-value < 0.05.

**RESULTS**

Metagenome assemblies

Analysis of the metagenome data identified 36 archaeal, 138 bacterial (Table S14), and 5 eukaryotic (**Table S13**) reference MAGs. Of the prokaryotic MAGs, which were dereplicated at 98% ANI (strain level), 40 had high (>90% completeness and <5% contamination), 109 had medium (≥50% and <10%), and 25 had low (<50% and <10%) completeness. Whereas a taxonomically broad range of bacterial MAGs were identified, 66 (47.83%) were classified as Actinobacteriota, 27 (19.57%) as Proteobacteria, and the remaining 45 were one of 16 different phyla. Of the 36 archaeal MAGs, 20 were Thermoplasmatota, 9 were Thermoproteota, 6 were Micrarchaeota, and 1 was Nanoarchaeota. The abundance and diversity of viruses in these metagenome data are described by Benites et al. 2024 ^45^.

Of the five eukaryotic MAGs, one was classified as the amoeba *Acanthamoeba* (57.7% complete using the BUSCO eukaryota_odb10 datasets), one was from the fungal genus *Acidomyces* (93.1% complete, capnodiales_odb10), one was from the fly order Diptera (23.7% complete, diptera_odb10), one was of the ciliate genus *Stylonychia* (83.1% complete, alveolata_odb10), and the last was from the green algal class Trebouxiophyceae (57.9% complete, chlorophyta_odb10) (**Table S13**).

Network analysis of the MAG data

Four major hubs of co-abundant MAGs were present in the network that was built. One of these hubs (green circle in **Fig. S1**) contains taxa with high representation in the creek biofilm samples (including *C. merolae*), one of the hubs (red circle) represents MAGs with high representation in the soil and some of the endolithic (mostly replicate 4) samples (including *G. yellowstonensis* and the other eukaryotes), with the two remaining hubs (orange and purple circles) being MAGs with high representation in either replicate 4 or replicates 1-3 (respectively) of the endolithic samples. Interestingly, MAGs in the creek biofilm hub show a tighter correlation than do the soil or endolithic samples, suggesting that this habitat, which is less species-rich, has a more stable assemblage of organisms than do the other, more heterogeneous soil and endolithic habitats.

SNP analysis of Lemonade Creek Cyanidiophyceae populations

For *C. merolae*, enriched GO terms with the highest *p*-value for genes in the top 5% block (20 kb size, 2 kb step) were related to the meiotic cell cycle in the Soil population, whereas in the Creek biofilm population, GO terms related to secondary alcohol biosynthetic and metabolic process were highly enriched. The Endolithic population showed chloride transport and sensory perception of chemical stimuli GO terms as highly enriched (**Table S20**). For *G. yellowstonensis*, enriched GO terms with the highest *p*-value for genes in the top 5% block (20 kb size, 2 kb step) were related to the ribosome in the Creek population, whereas the Endolithic and Soil populations showed only a few enriched GO terms, such as tissue homeostasis. Because of the low number of SNPs in the creek biofilm data, we did this analysis using unfiltered SNP data from *G. yellowstonensis* (**Table S20**).

Cyanidiophyceae environment SNP analysis

For *C. merolae*, enriched GO terms with the highest *p*-value for genes in the top 5% block (20 kb size, 2 kb step) were related to the meiotic cell cycle in the soil population, whereas in the creek biofilm population, GO terms related to secondary alcohol biosynthetic and metabolic process were highly enriched. The endolithic population showed chloride transport and sensory perception of chemical stimuli GO terms as highly enriched (**Table S20**). We did not do this analysis for *G. yellowstonensis* because of the low number of SNPs in the creek biofilm data. Therefore, we only generated GO terms from the comparison between the endolithic and soil populations and found that terms associated with DNA unwinding involved in DNA replication and mitochondrial RNA metabolic process were significantly enriched (**Table S20**).

Cell cycle and the diurnal phototrophy-heterotrophy cycle

Light responses, the cell cycle, and metabolism of *C. merolae* and *G. yellowstonensis* in the Lemonade Creek habitats were evaluated using the poly-A expression pattern of target genes (**Table S21**). *G. yellowstonensis* was evaluated based solely on soil data because of the paucity of reads from the creek biofilm samples (**Fig. 2A**). In *C. merolae*, two copies of the cryptochrome DASH gene, a subclade of the cryptochrome/photolyase family, some of which accumulate at midday in other organisms ^50^, exhibited a diurnal pattern, peaking at midday in the creek samples (**Fig. S9A**). In *C. merolae*, as observed in lab cultures over the diurnal cycle (12-hour light/12-hour dark) ^51^, genes involved in glycolysis/gluconeogenesis (e.g., 6-phosphofructokinase; **Fig. S9C**), the entrance to the TCA cycle (e.g., dihydrolipoamide dehydrogenase; E3 component of pyruvate dehydrogenase complex; **Fig. S9E**), and the antenna of the photosystem (chlorophyll *a* binding protein; **Fig. S9G**) were upregulated during the day and downregulated at night. Conversely, cell division genes (S- and M-phase cyclins and the chloroplast division gene *ftsZ*) and lactate fermentation (L-lactate dehydrogenase) were upregulated during the night in the creek samples (**Fig. S9E**). These results suggest that, in *C. merolae*, as observed in lab cultures ^52^, in the creek environment where this alga coexists with other microorganisms, the cells exhibit higher respiratory activity due to photosynthesis during the day but lower respiratory activity at night, with an increased reliance on fermentation for ATP production. However, in the soil samples, these diurnal rhythms were not evident in *C. merolae* and *G. yellowstonensis* (**Fig. S9**). We postulate that this is explained by the cells in the soil being exposed to lower light levels during the day and due to the mixotrophic lifestyle of *G. yellowstonensis*.

Arsenic and mercury detoxification in Lemonade Creek

We investigated RiboMinus RNA-seq data from the creek biofilm and soil habitats to understand mercury and arsenic detoxification in these YNP habitats. Orthogroups (OGs) were constructed using all Cyanidiophyceae reference genomes plus reference MAGs from the metagenome data. Whereas OG clustering reconstituted the expected gene families in each Cyanidiophyceae genome, there were some notable differences. The local *C. merolae* and *G. yellowstonensis* contained more *arsM* (2 and 2 genes, respectively; OG0000651) and *merA* (4 and 3 genes, respectively; OG0000025) than in the existing genome data (**Fig. 3A; Table S18**). This suggests these OGs include related genes with similar functions, or that the gene families are larger in wild populations than previously estimated.

Orthogroup (OG) analysis regroups some of the ArsA, ArsB, “arsenical pump-driving ATPase”, and “arsenic transporter” genes together into combined OGs. OG0000997 and OG0000581 comprise genes annotated as “arsenic transporter”, and ArsA or ArsB (respectively). Both OGs include sequences from prokaryotes and other eukaryotes in Lemonade Creek. Two of the new OGs, OG0006594 and OG0005263, comprise only genes from *Galdieria* annotated as ArsA or “arsenic transporter” (respectively). The *arsC* genes were split into two OGs, one (OG0000808) that is broadly shared with prokaryotes, and another (OG0024314) that is specific to two *Galdieria* isolates (Gsulp_MtSh and Gyell_5587_1; **Table S19**). The *arsH* genes form a single OG with nearly the same composition as in the genome data (**Fig. 3A**). Only four genes in the *arsH* OG are from prokaryotes, and one from the acidophilic fungus *Acidomyces*. There were no viral sequences identified in any of these OGs, suggesting that Lemonade Creek viruses do not carry these detoxification genes.

An As(III) efflux pump is encoded by *arsB,* which only occurs in *G. yellowstonensis* 5587.1 (two copies). The *arsA* gene encodes an ATPase that is present in *G. yellowstonensis* 5587.1 and *C. merolae* 10D (two copies). ArsA forms a complex with ArsB, providing ATP-derived energy for ArsB to actively extrude As(III) from the cells ^52^. The *arsM* genes encode arsenic methyltransferase ^53^ and are present in *G.* *yellowstonensis* 5587.1 (one copy) and *C. merolae* 10D (two copies). ArsM methylates As(III), and depending on the specific enzyme, results in variously methylated arsenicals, ranging from the highly toxic methyarsenite (MAs(III)) to trimethylarsine, which is volatile and leaves the cell (**Fig. 3B**). Therefore, ArsM can serve as an arsenic detoxification mechanism ^54^, but also has an arsenic activation mechanism. Both functions could potentially influence the composition of closely associated cells in the mat community. *ArsM*-expressing *E. coli* cells exhibit As(III) resistance ^55^ while killing co-cultured bacterial cells in the presence of As(III) ^54^. Two ArsM encoding genes cloned from *C. merolae* 5508 have robust As(III) methyltransferase activity and serve to detoxify As(III) ^55^. Given the high sequence similarity (>50% identity) with the active ArsM homologs from *C. merolae* 5508, it is highly likely that Cyanidiophyceae ArsM function as As(III) methyltransferases. The *arsH* genes encode an NADPH-dependent FMN oxidoreductase that oxidizes highly toxic MAs(III) to inert MAs(V), and thus confers MAs(III) resistance ^56^. In Cyanidiophyceae that inhabit Lemonade Creek, *arsH* is only found in *G. yellowstonensis* (two copies) and shares similarity with bacterial *arsH*. Theoretically this could function as a defense mechanism against MAs(III), but also in pathways unrelated to arsenic transformation ^57^.

Other metal resistance genes in the Lemonade Creek data

Beyond *ars* and *mer*, we identified genes in both YNP algal species that confer resistance to copper (Cu), zinc (Zn), and manganese (Mn) (**Fig. S10**). In the case of copper, genes encoding HMA1 ^58^ and HMA5 ^59^ that import Cu^+^ into the plastid and cytoplasm (respectively) are present. However, genes for Cu export from the cytoplasm were absent. Only the NRAMP family genes for Mn^2+^ import was identified in both algal species. ZIP proteins import Zn^2+^, which is exported by ZnTs. Gene encoding ZIP and ZnT for cytoplasmic import were present in both species, however, proteins for Golgi import were restricted to *C. merolae*. Because the Golgi can store Zn^2+^ and harbor various Zn metalloenzymes, the loss of these genes may make *G. yellowstonensis* more susceptible to elevated Zn levels in the environment. The expression patterns of genes involved in Cu, Zn, and Mn are the same as *mer,* suggesting that metal detoxification follows the diurnal cycle.

The integrated HGT model

Given the strong partitioning of arsenic detoxification uncovered at Lemonade Creek, we asked whether *ars* genes of HGT origin in Cyanidiophyceae that are not highly expressed may have been freed from selective constraints and gained novel functions. This outcome is predicted by the integrated HGT model (IHM) for eukaryotes which posits that when conditions that initially favor HGT fixation diminish or disappear (e.g., due to toxin absence or the evolution of community detoxification), foreign gene(s) may survive if they are integrated into a broader stress (or other) responses that favor retention. The IHM is akin to the Black Queen Hypothesis, whereby community interactions (e.g., provision of public goods [metabolites]) can drive prokaryote genome reduction ^60^. The IHM contrasts with the standard model of HGT, as for *mer* genes, whereby this gene has retained its original function for hundreds of millions of years ^57^. Although we lack functional data to test this idea with Cyanidiophyceae *ars* genes, inspection of gene co-expression data for the 5 *arsH* genes in *G. partita* SAG21 is consistent with the IHM, showing that they are strongly linked with modules dominated by protein translation and photosynthesis. Furthermore, some ArsH isoforms in *G. partita* SAG 21 have lost the conserved binding motifs for the phosphate group of flavin mononucleotide (FMN) and nicotinamide adenine dinucleotide phosphate (NADP^+^) ^57^; see main text (**Fig. 4**).

Other metabolites of interest found at YNP

Several other metabolites may influence population dynamics at Lemonade Creek. Phytosphingosine (PSN), a long-chain amino alcohol, is a major toxic metabolite in cyanobacterial blooms, where it induces cell apoptosis by disrupting the mitochondrial membrane ^61^] (**Fig. 5**). We found that PSN was highly accumulated at TP3 and TP2, in the supernatant of the creek biofilm and in the soil samples, respectively. However, intracellular PSN is involved in structural integrity and in mediating cellular processes ^62^. Creatine is a non-proteogenic amino acid that can accelerate the circadian clock of algae and may act as inhibitor of bacterial replication ^63^. The peak of creatine abundance coincided with the peak of algal growth in the creek biofilm, whereas guanidineacetic acid, a precursor of creatine, was elevated extracellularly at TP2 in the creek and soil. Finally, the presence of different forms of glycerol can denote restructuring of the community vis-à-vis lipids or can suggest a trophic chain between bacteria or fungi as producers, and red algae, equipped with the necessary enzymatic arsenal, as consumers ^64^. The rapid decline in abundance at TP4 in the soil and the moderate overnight decline in the creek correlate with algal growth patterns (poly-A RNA data). Conversely, the pattern between the extracellular N-acetyl-D-glucosamine (peak at TP2) and intracellular N-acetyl-glucosaminyl-asparagine (peak at TP3) may indicate the salvage of degraded cell wall components originating from other bacteria or fungi.

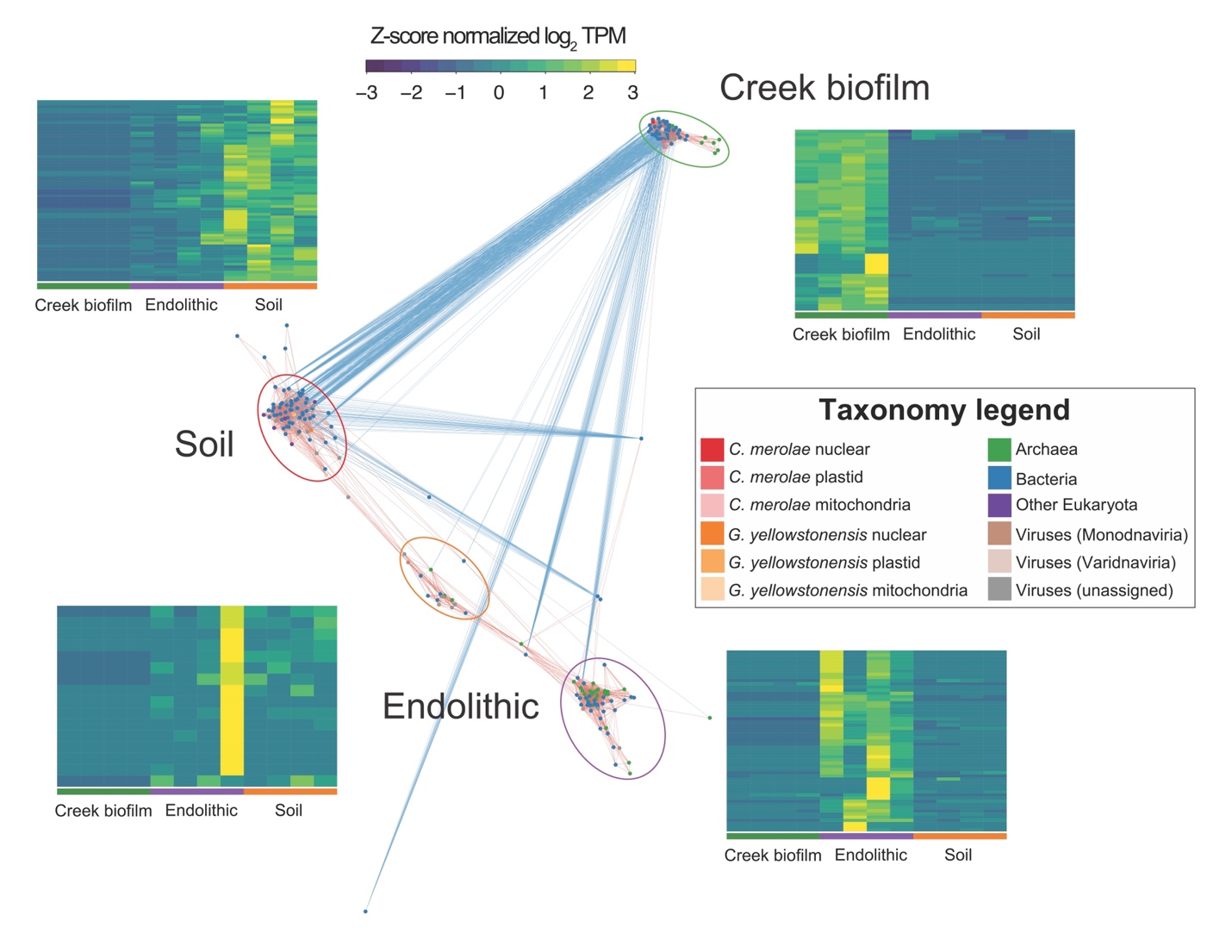

**Fig. S1. Co-abundance network of Yellowstone reference MAGs.** Co-abundance was estimated using the TPM normalized values from the mapped metagenome samples, with the data scaled using z-score normalization before adjacency calculation. Nodes are colored according to the legend at the bottom of **Fig. 1B**, and the edges are colored such that nodes with positive correlation are red and those with negative correlation are blue. Each of the major hubs of nodes are highlighted using colored circles and the expression patterns of the MAGs in each of these hubs are shown in the adjacent heatmaps. The colors used in the heatmap are shown at the top of the image.

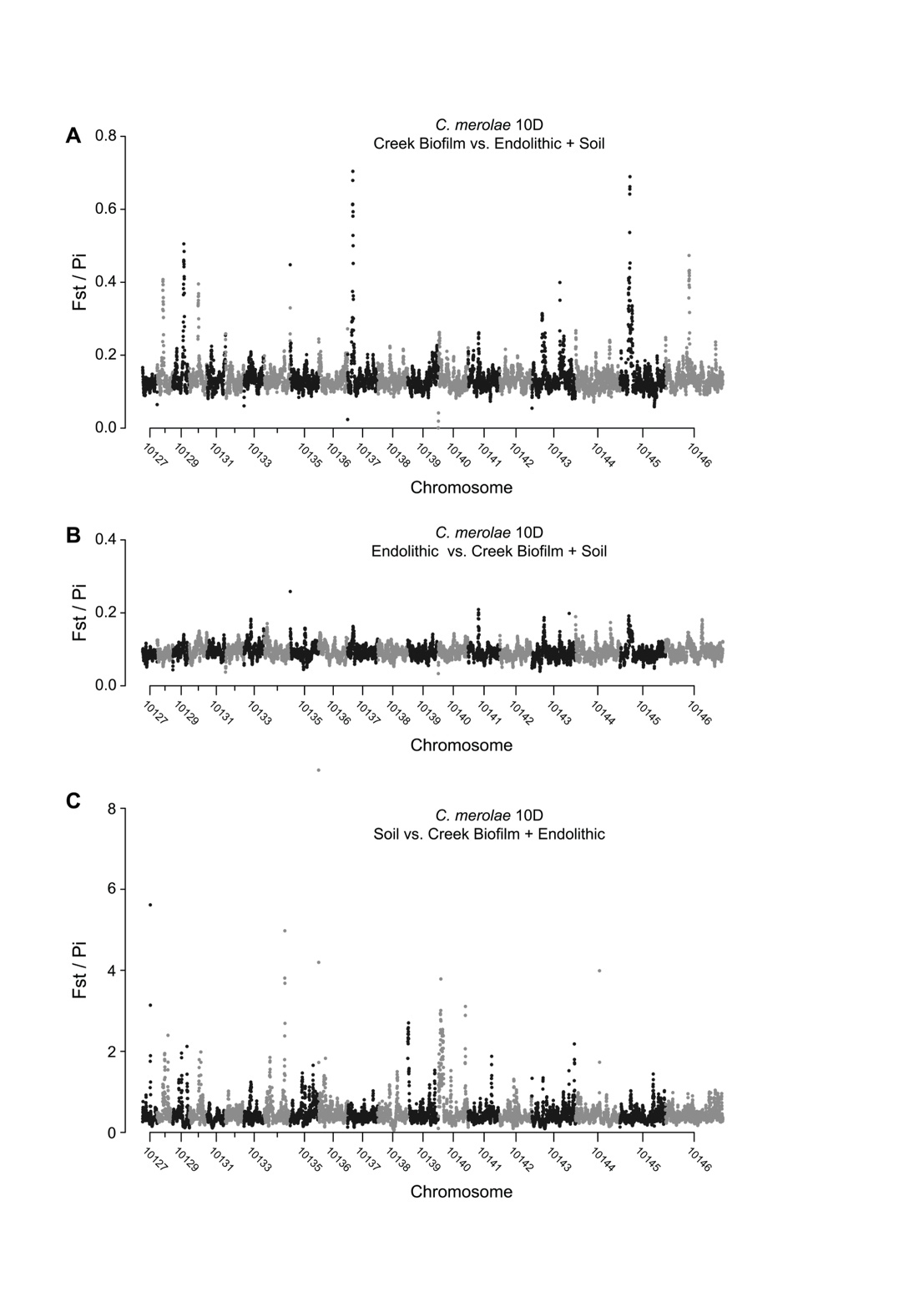

**Fig. S2. Fixation index (F_ST_) divided by nucleotide diversity (Pi) of three populations using the reference genome of *C. merolae* 10D.** Because a high F_ST_ value (range from 0 to 1) implies that populations under study are strongly separated, and a low Pi (average pairwise difference between all possible pairs of individuals in the studied sample) indicates the region is overall less variable, F_ST_ was divided by Pi (window size 20 kb, window step 2 kb) to identify regions putatively undergoing selective sweeps in the Creek biofilm (A), Endolithic (B), and Soil (C) populations. Each population shows a distinct pattern, indicating different selective pressures in each habitat.

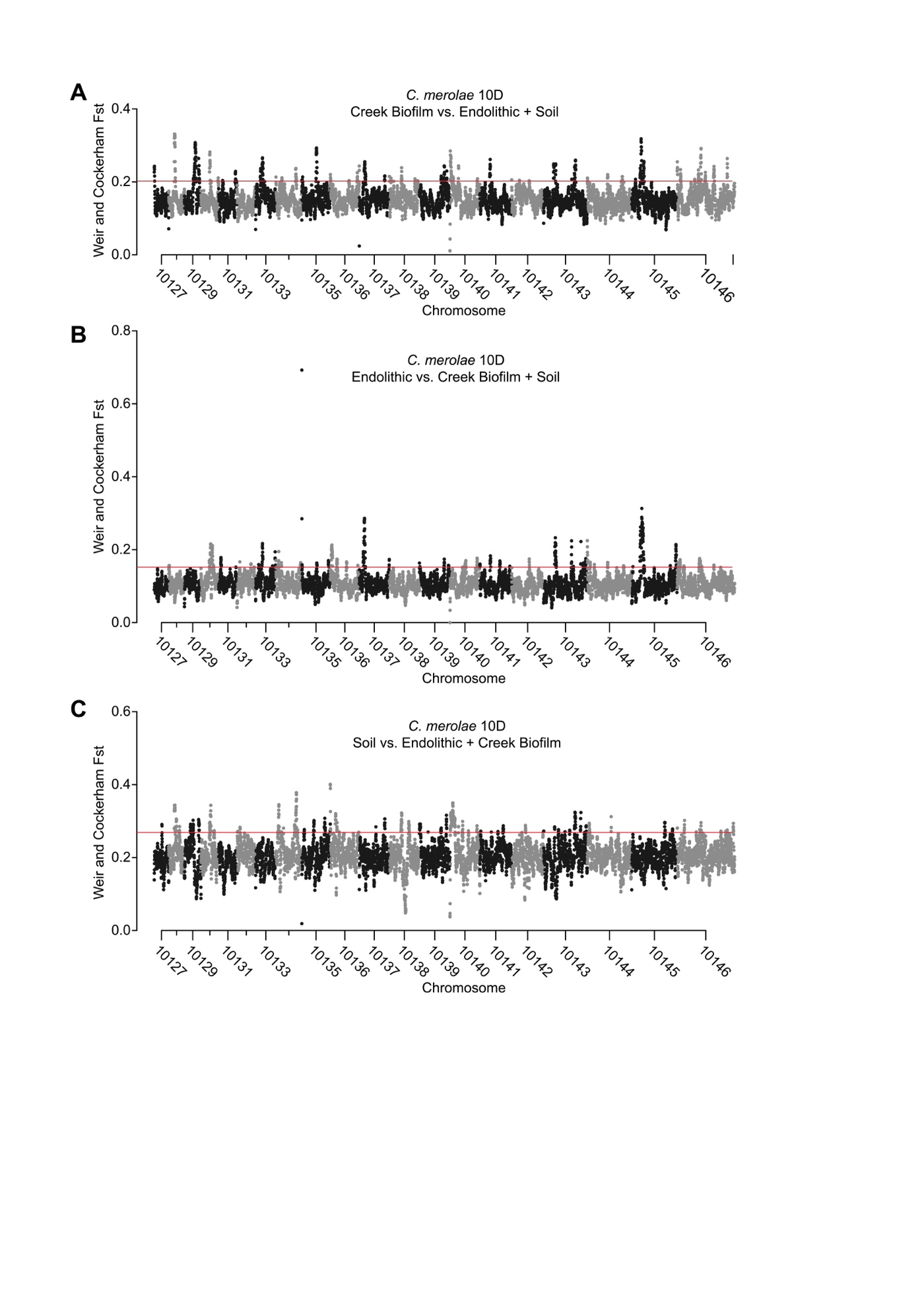

**Fig. S3. F_ST_ of three populations using the reference genome of *C. merolae* 10D.** Weir and Cockerham F_ST_ values for the Creek biofilm (A), Endolithic (B), and Soil (C) populations were calculated using a 20 kb window size (step size 2 kb). Each population shows distinct patterns. The red lines indicate top 5% F_ST_ value (Creek biofilm, 0.20; Endolithic, 0.15; Soil, 0.27).

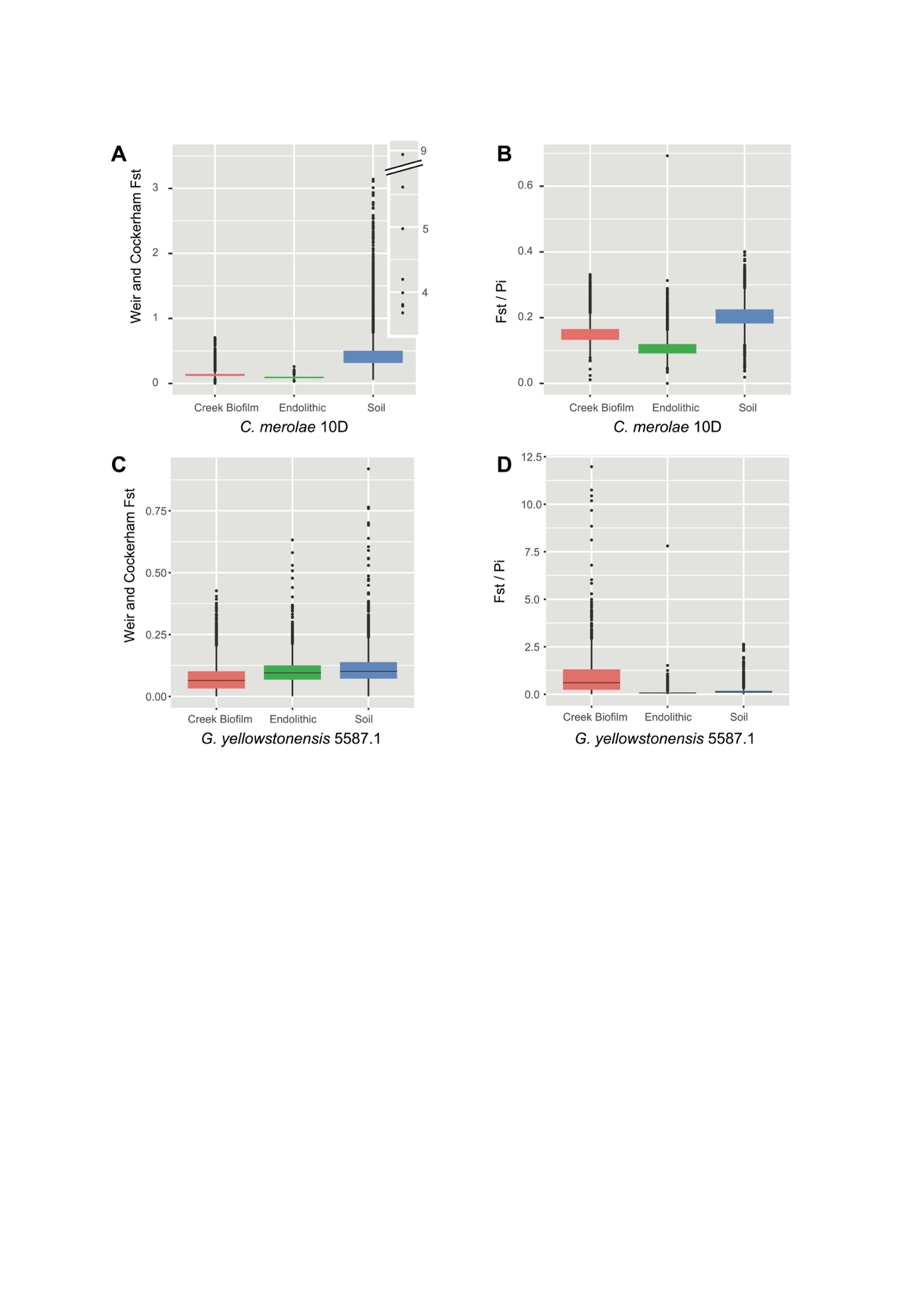

**Fig. S4. Boxplot of F_ST_ and F_ST_/Pi values of three YNP populations using the reference genomes of *C. merolae* 10D and *G. yellowstonensis* 5587.1.** The box includes the 25th (Q1) to 75th (Q3) percentiles of the data with the median value in a thick line. Upper and lower whiskers indicate values within 1.5 times interquartile range (Q3 – Q1) above Q3 and below Q1, respectively. F_ST_ and F_ST_/Pi value of the Creek biofilm, Endolithic, and Soil populations of *C. merolae* 10D (A, B). Soil populations show significantly higher values than the other populations. F_ST_ and F_ST_/Pi values of the Creek biofilm, Endolithic, and Soil populations of *G. yellowstonensis* 5587.1 (C, D). For *G. yellowstonensis* 5587.1, calculations were done based on the unfiltered SNP data, due to an insufficient number of SNPs from the Creek biofilm population.

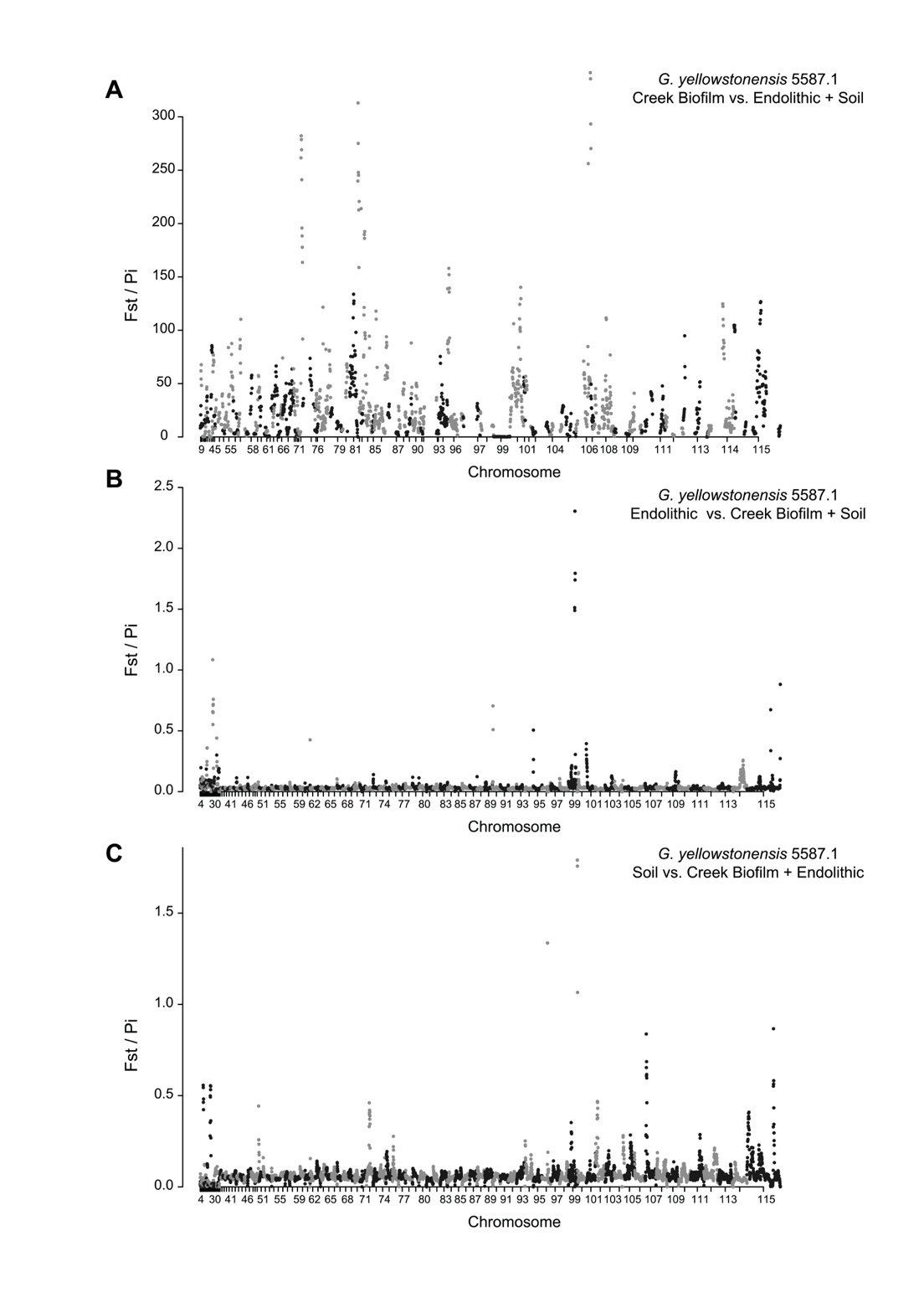

**Fig. S5. Fixation index (F_ST_) divided by nucleotide diversity (Pi) of three populations using the reference genome of *G. yellowstonensis* 5587.1.** As in **Fig. S2**, F_ST_ was divided by Pi (window size 20 kb, window step 2 kb) to infer regions with a putative selective sweep in the Creek biofilm (A), Endolithic (B), and Soil (C) populations. Each population shows a distinct pattern, indicating different selective pressures in each habitat. Because of the low SNP number from the Creek biofilm population, low Pi resulted in very high values in panel (A). Calculations were done based on the unfiltered SNP data, due to an insufficient number of SNPs from the Creek biofilm population.

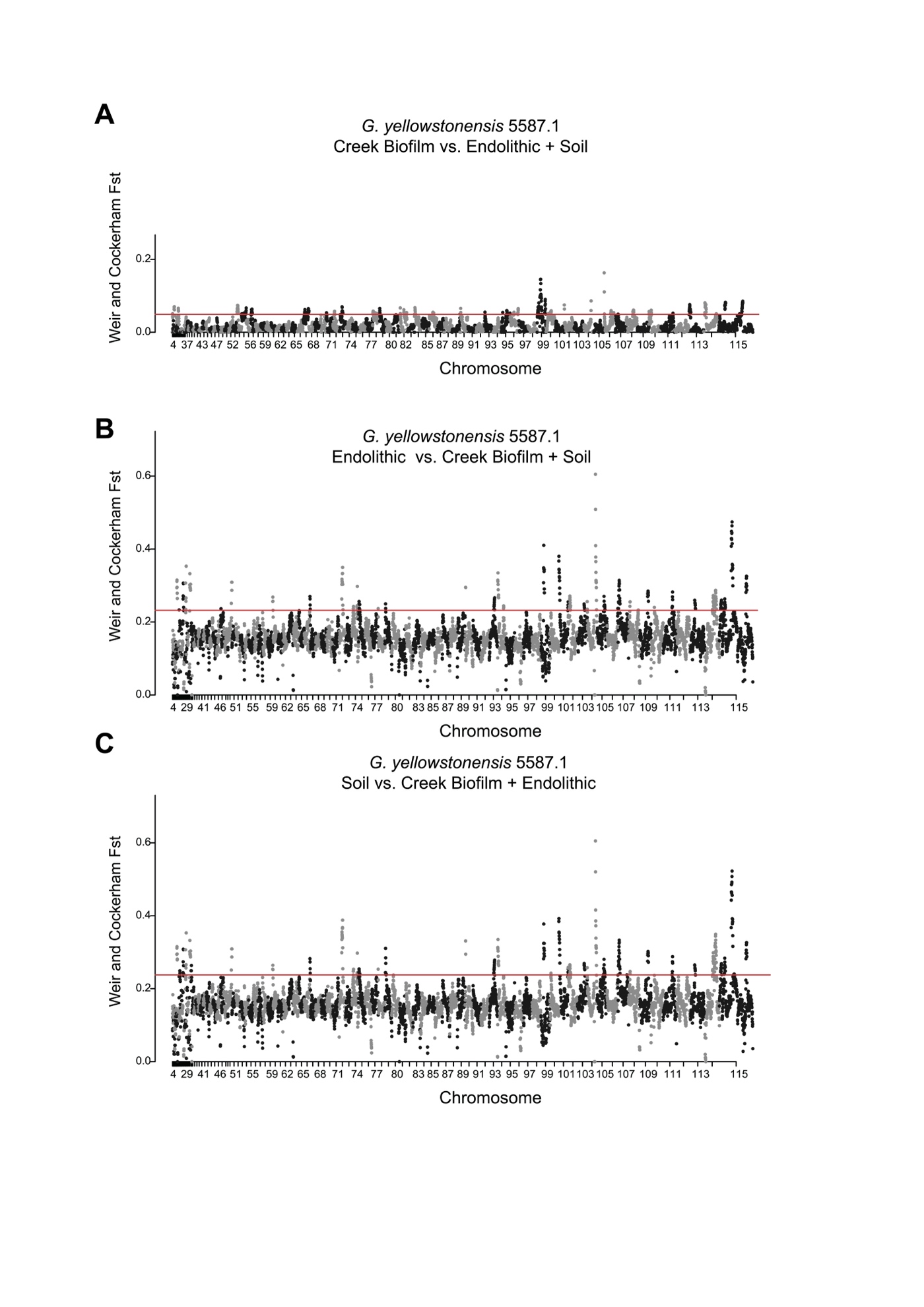

**Fig. S6. F_ST_ of three populations using the reference genome of *G. yellowstonensis* 5587.1.** Weir and Cockerham F_ST_ values for the Creek biofilm (A), Endolithic (B), and Soil (C) populations were calculated within 20 kb window size (step size 2 kb). The red lines indicate the top 5% F_ST_ value (Creek biofilm, 0.49; Endolithic, 0.23; Soil, 0.24). The Creek biofilm population shows a different pattern, whereas the Endolithic and Soil populations show similar patterns. Calculations were done based on unfiltered SNP data, due to an insufficient SNP number from the Creek biofilm population.

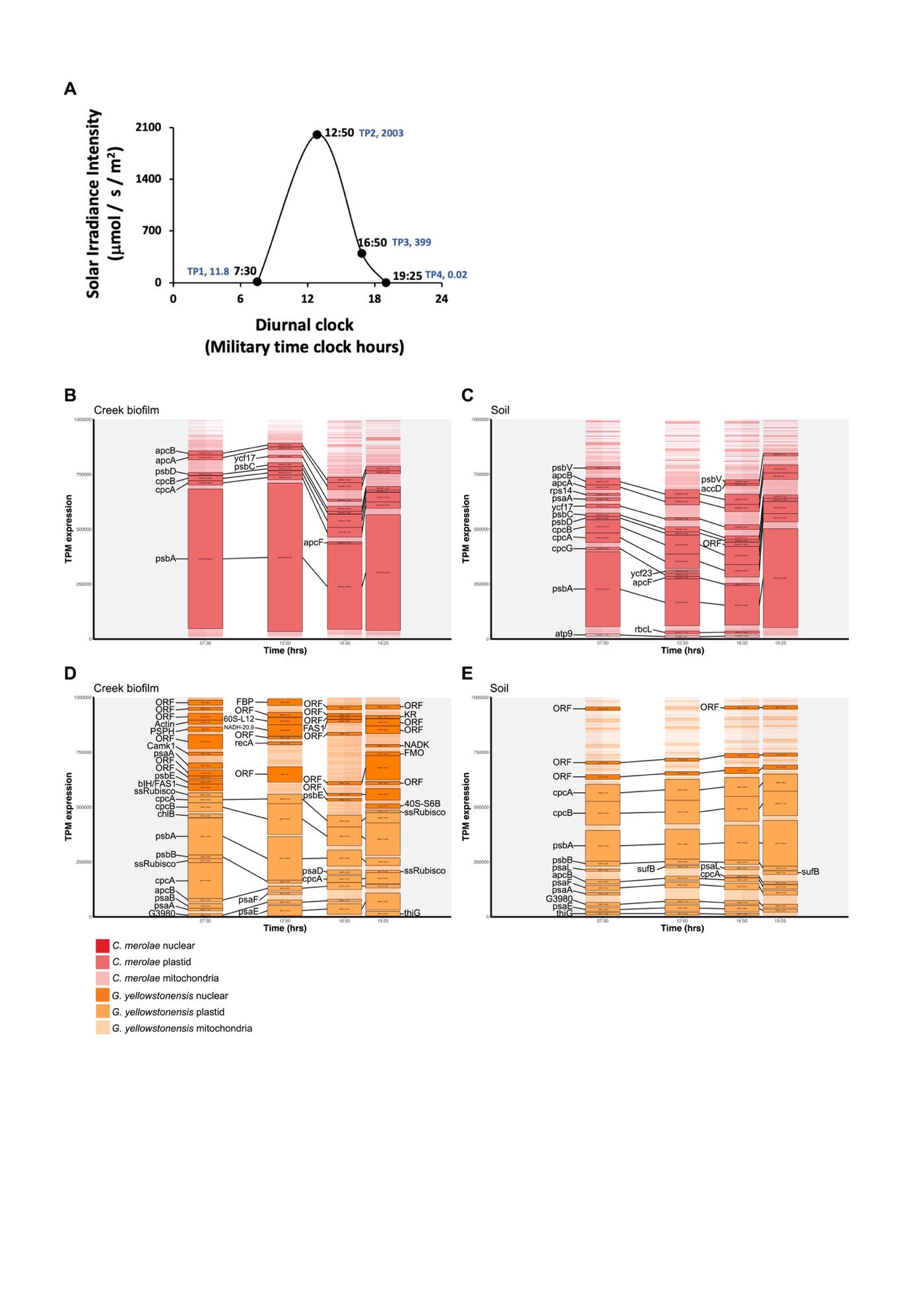

**Fig. S7. Light levels at YNP and per species TPM values of major expressed genes.** (A) Surface light levels measured at Lemonade Creek, YNP on October 10, 2021. The sampling times, sample numbers, and irradiance levels are indicated. The peak irradiance level is 2,003 mmol m^-2^ s^-1^ at midday, TP2. RiboMinus RNA-seq readers that mapped to genes from the (B, C) *C. merolae* or (D, E) *G. yellowstonensis* 5587.1 nuclear and organelle reference genomes were extracted and used to recalculate TPM values for the respective gene sets. Genes with > 1% total abundance (a TPM > 10,000) in each dataset at each time point are highlighted, have the percentage of total TPM shown in brackets at each time point, and the gene symbols of their putative functions displayed along their side. Genes highlighted across multiple time points are connected by black lines. A legend describing the colors used is shown at the bottom of the figure. Gene symbol abbreviations: *apcB*: allophycocyanin beta chain; *apcA*: allophycocyanin alpha chain; *psbD*: photosystem II D2 protein; *cpcB*: phycocyanin beta chain; *cpcA*: phycocyanin alpha chain; *psbA*: photosystem II D1 (Q[b]) protein; *ycf17*: ELIP-like protein; *psbC*: photosystem II 44 kDa apoprotein; *apcF*: allophycocyanin B18 chain; *psbV*: cytochrome c550; *rps14*: 30S ribosomal protein S14; *psaA*: Photosystem I P700 chlorophyll a apoprotein A1; *cpcG*: phycobilisome rod-core linker polypeptide; *atp9*: ATP synthase F0 subunit 9; *ycf23*: ycf23; *apcF*: allophycocyanin B18 chain; *rbcL*: ribulose-1,5-bisphosphate carboxylase/oxygenase large subunit; *accD*: acetyl-CoA carboxylase carboxyl transferase beta; ORF: unnamed protein product/hypothetical protein; *PSPH*: phosphoserine phosphatase; *Camk1*: Calcium/calmodulin-dependent protein kinase type 1; *psbE*: photosystem II cytochrome b559 alpha subunit; bIH/FAS1: beta-Ig-H3/fasciclin; ssRubisco: ribulose-1,5-bisphosphate carboxylase/oxygenase small subunit; *chlB*: protochlorophyllide reductase ChlB subunit; *psbB*: photosystem II 47 kDa protein; *psaB*: photosystem I P700 chlorophyll a apoprotein A2; G3980: putative photosynthetic reaction centre, L/M, Photosystem antenna protein-like protein; *FBP*: fructose-bisphosphate aldolase; 60S-L12: 60S ribosomal protein L12; NADH-20.9: NADH-ubiquinone oxidoreductase 20.9 kDa subunit; *recA*: recombinase RecA; *psaE*: photosystem I reaction center subunit IV; *FAS1*: Uncaracterized surface protein containing fasciclin (FAS1) repeats; *psaD*: photosystem I subunit II; *psaF*: photosystem I reaction center subunit III; KR: ketol-acid reductoisomerase; *FMO*: flavin-containing monooxygenase FMO GS-OX-like 4; *NADK*: NAD kinase isoform X3; 40S-S6B: 40S ribosomal protein S6-B; *thiG*: thiamin biosynthesis protein G; *psaL*: Photosystem I reaction center subunit XI; *sufB*: Iron-sulfur cluster assembly protein SufB.

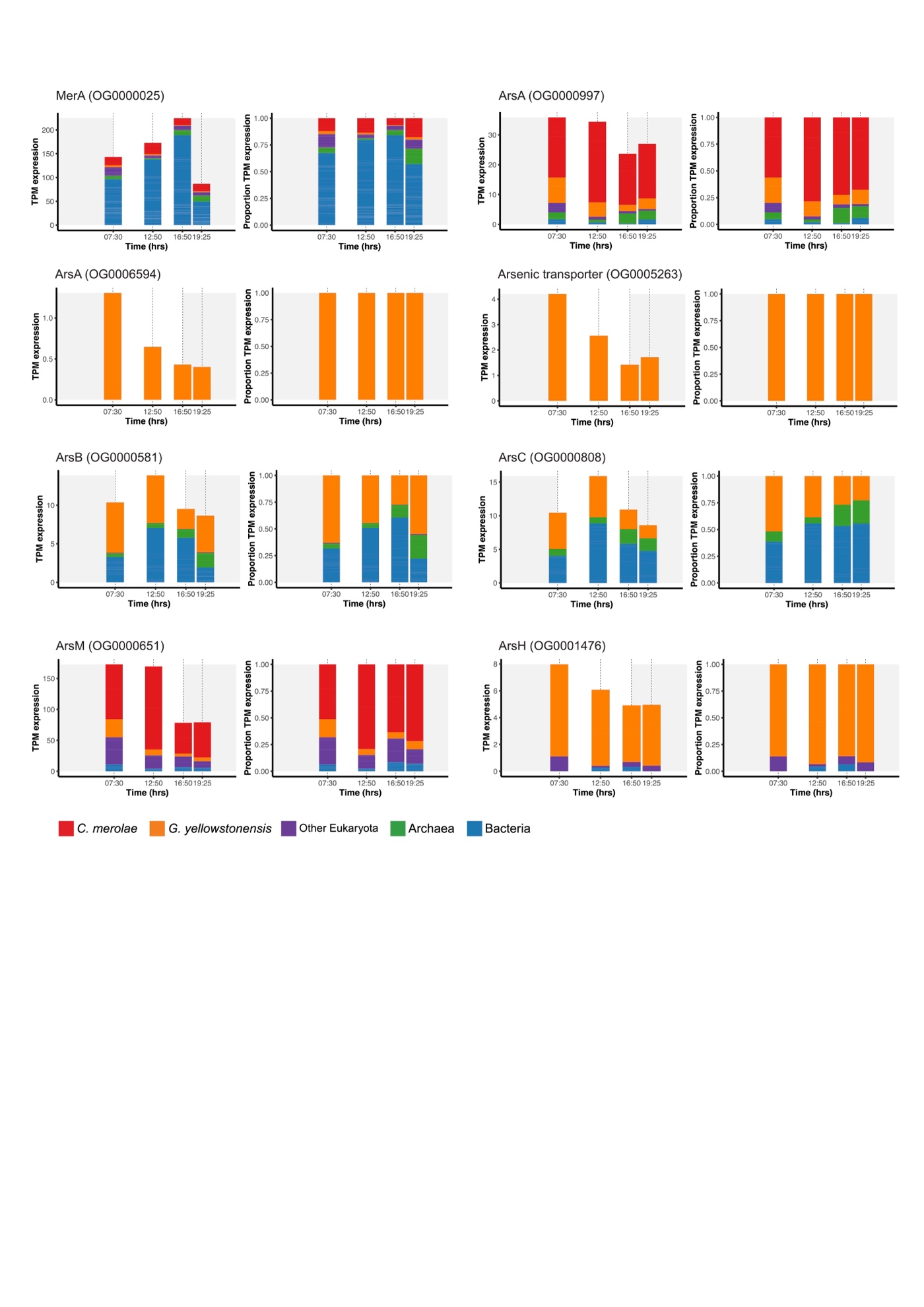

**Fig. S8. Analysis of *mer* and *ars* gene expression in the soil.** Contribution of taxonomic groups to the arsenic and mercury detoxification pathways in the Soil samples. The TPM expression values of all genes from an orthogroup identified as containing genes putatively from a specific step in the detoxification pathway is shown as a stacked bar graph (left). The proportion of TPM values contributed by each taxonomic group is shown as a stacked bar graph (right).

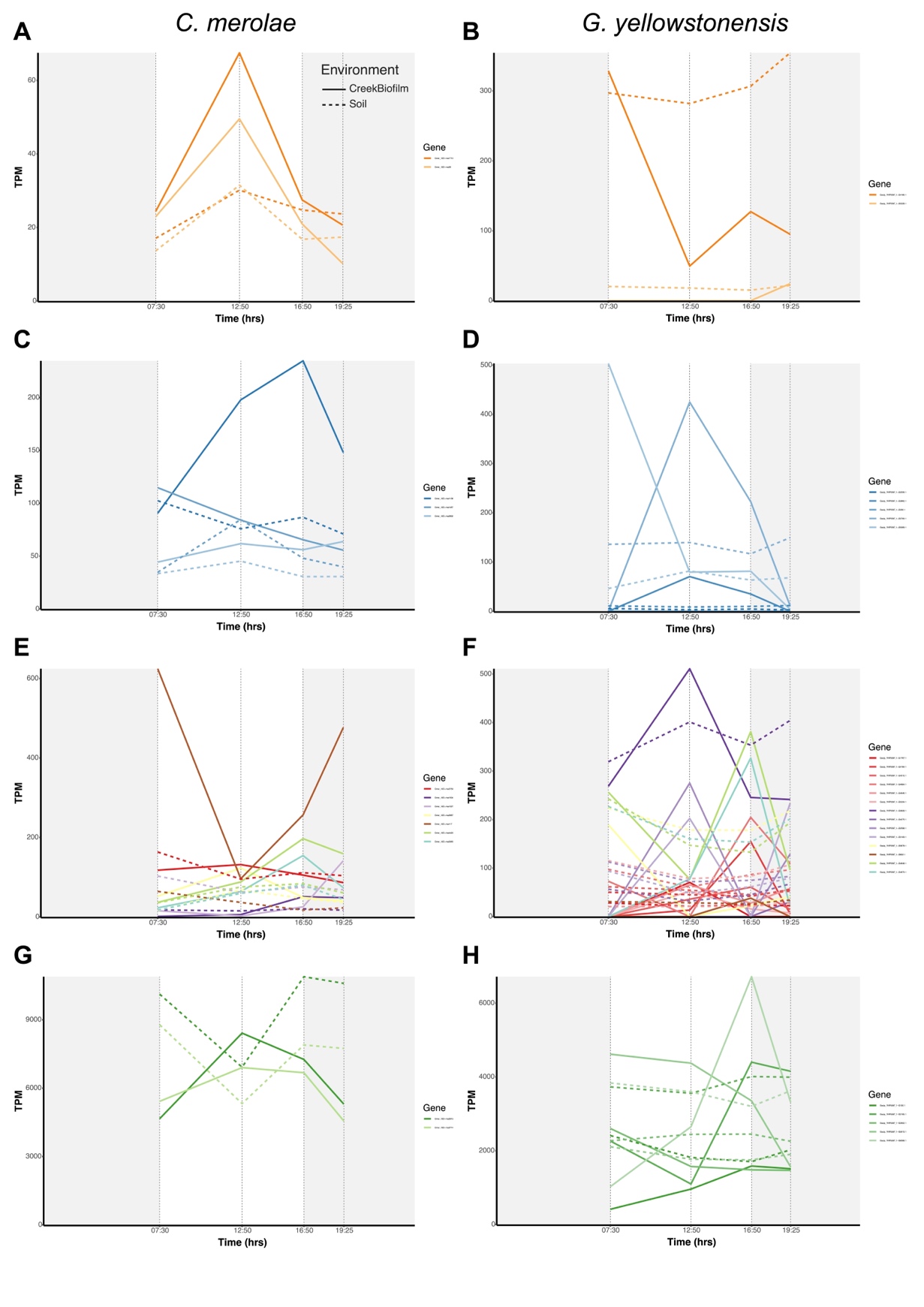

**Fig. S9. Expression of cell cycle associated genes.** TPM values calculated using Poly-A reads extracted from (A, C, E, G) *C. merolae* and (B, D, F, H) *G. yellowstonensis* nuclear and organelle genes. Gene names are shown on the right of each panel and have their putative functions described in **Table S21**. Expression of each gene in the Creek biofilm and Soil samples is displayed separately using solid or dashed lines (respectively).

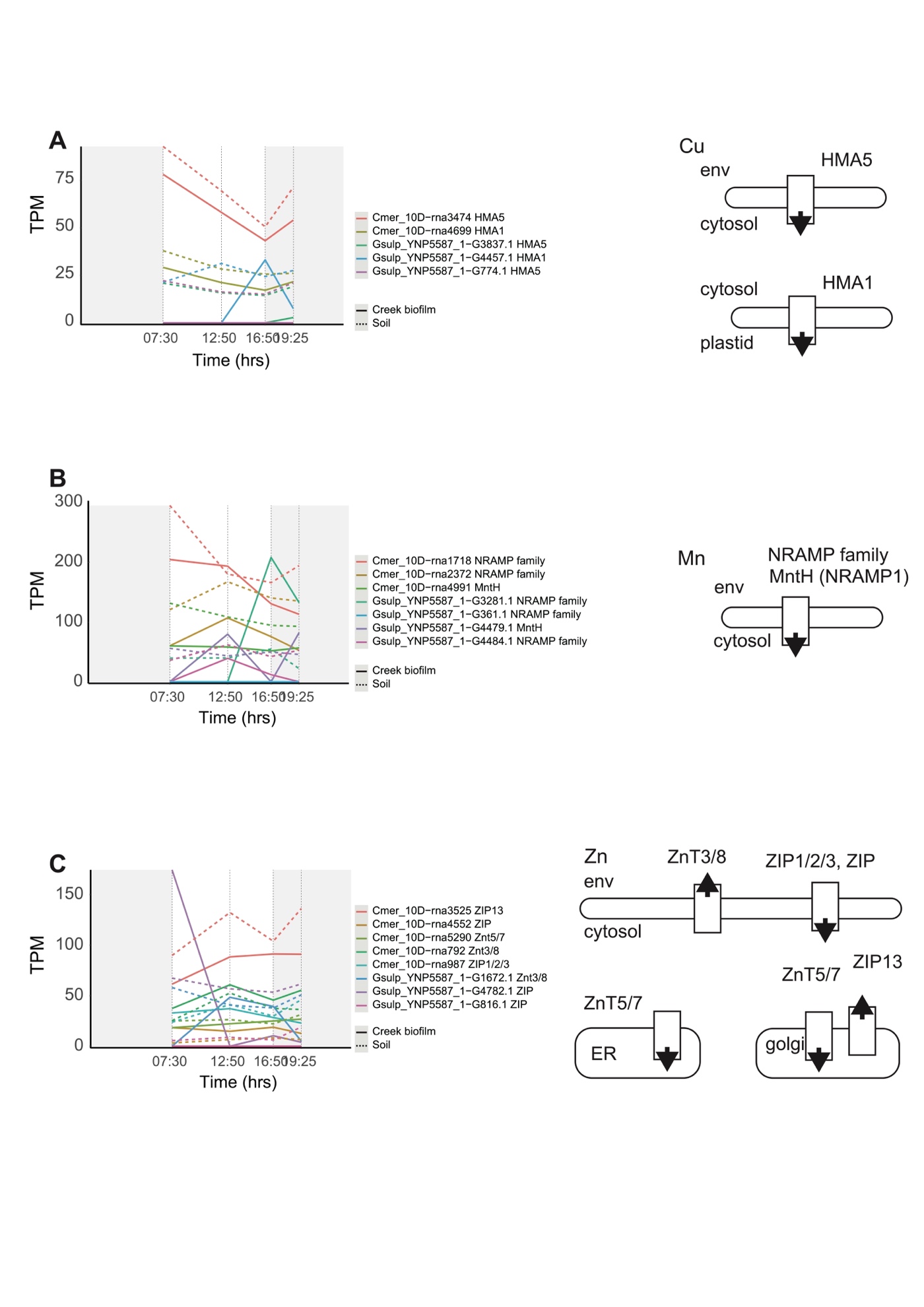

**Fig. S10. Expression of metal detoxification genes.** TPM values calculated using Poly-A reads extracted from *C. merolae* and *G. yellowstonensis* nuclear and organelle genes. The expression patterns of genes associated with (**A**) copper (Cu), (**B**) manganese (Mn), and (**C**) zinc (Zn) are shown along with diagrams of their putative functions (right side of each panel). Gene names are shown on the right of each panel with their putative functions.

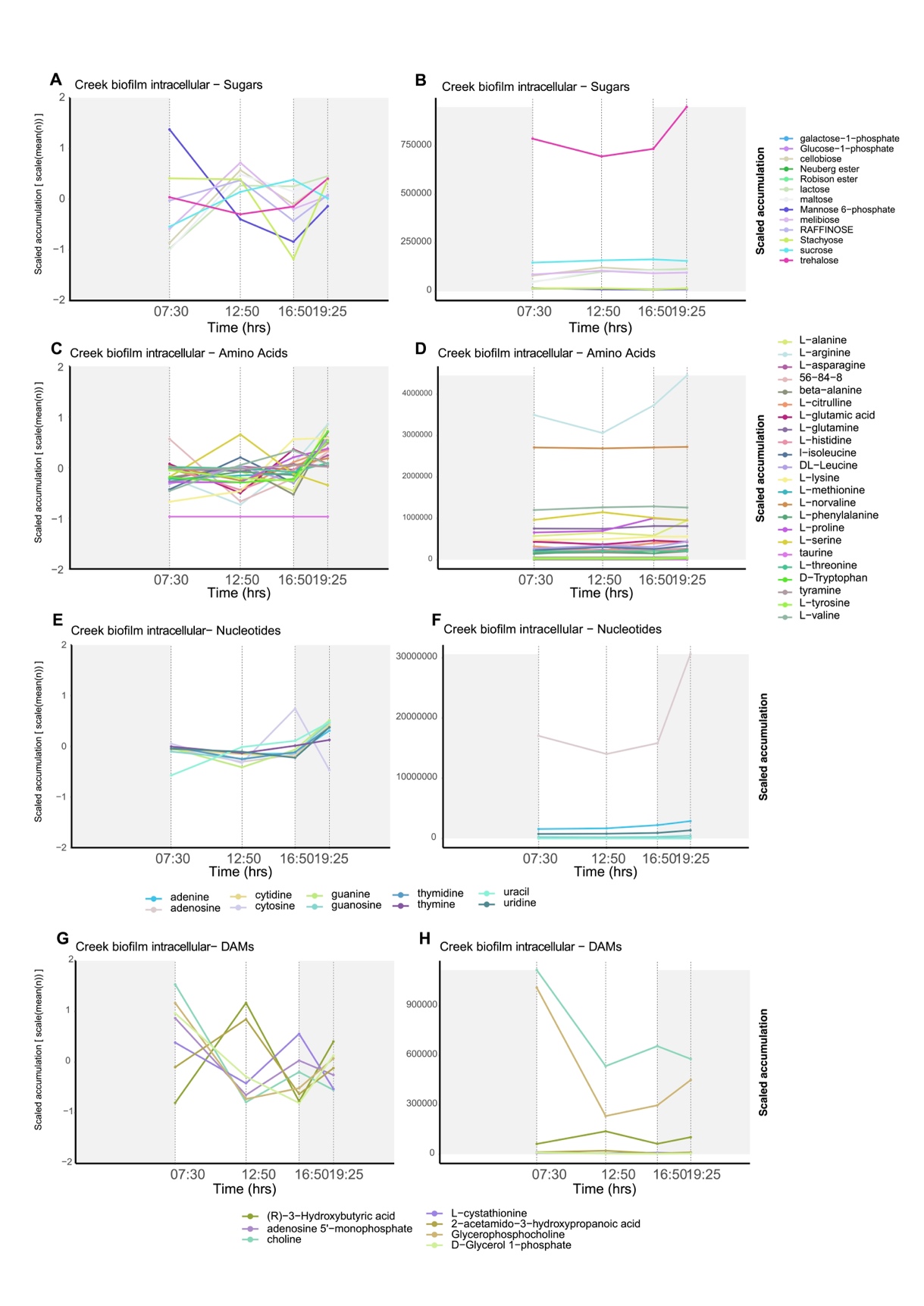

**Fig. S11. Accumulation patterns of metabolites in the targeted metabolome in the Creek biofilm intracellular samples.** The log_2_ z-score normalized and absolute intensity values of the sugar (A, B), amino acid (C, D), and nucleotide (E, F) metabolites identified in the targeted data are shown as line graphs. The same graphs are shown for the metabolites differentially accumulated (|FC| > 1 and *p*-value < 0.05) (DAMs) between any combination of samples (G, H). Each point in the line graph represents the average value of the n=4 replicates per time point sequenced. The colors used for each metabolite are shown in the legends.

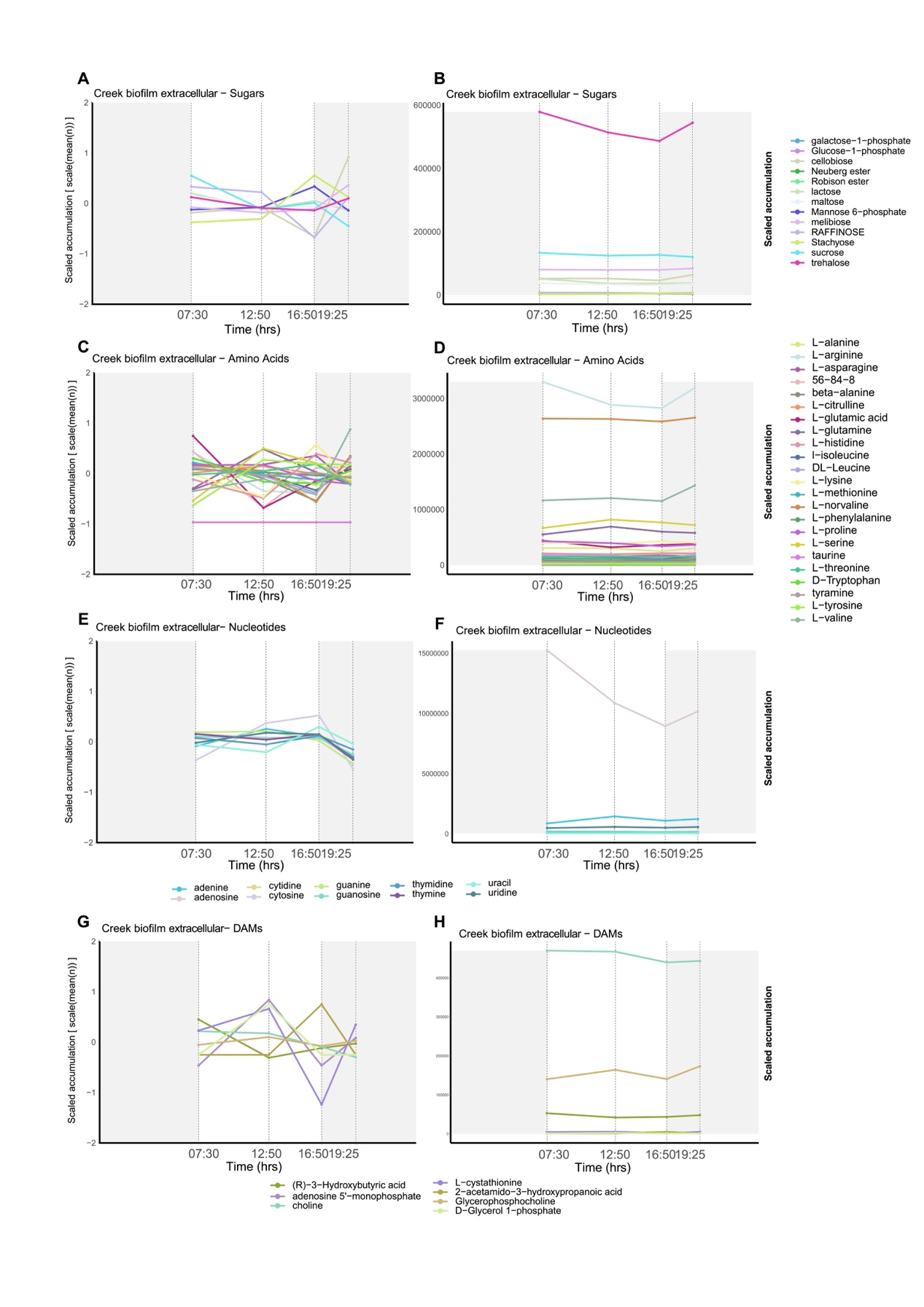

**Fig. S12. Accumulation patterns of metabolites in the targeted metabolome in the Creek biofilm extracellular samples.** The log_2_ z-score normalized and absolute intensity values of the sugar (A, B), amino acid (C, D), and nucleotide (E, F) metabolites identified in the targeted data are shown as line graphs. The same graphs are shown for the metabolites differentially accumulated (|FC| > 1 and *p*-value < 0.05) (DAMs) between any combination of samples (G, H). Each point in the line graph represents the average value of the n=4 replicates per time point sequenced. The colors used for each metabolite are shown in the legends.

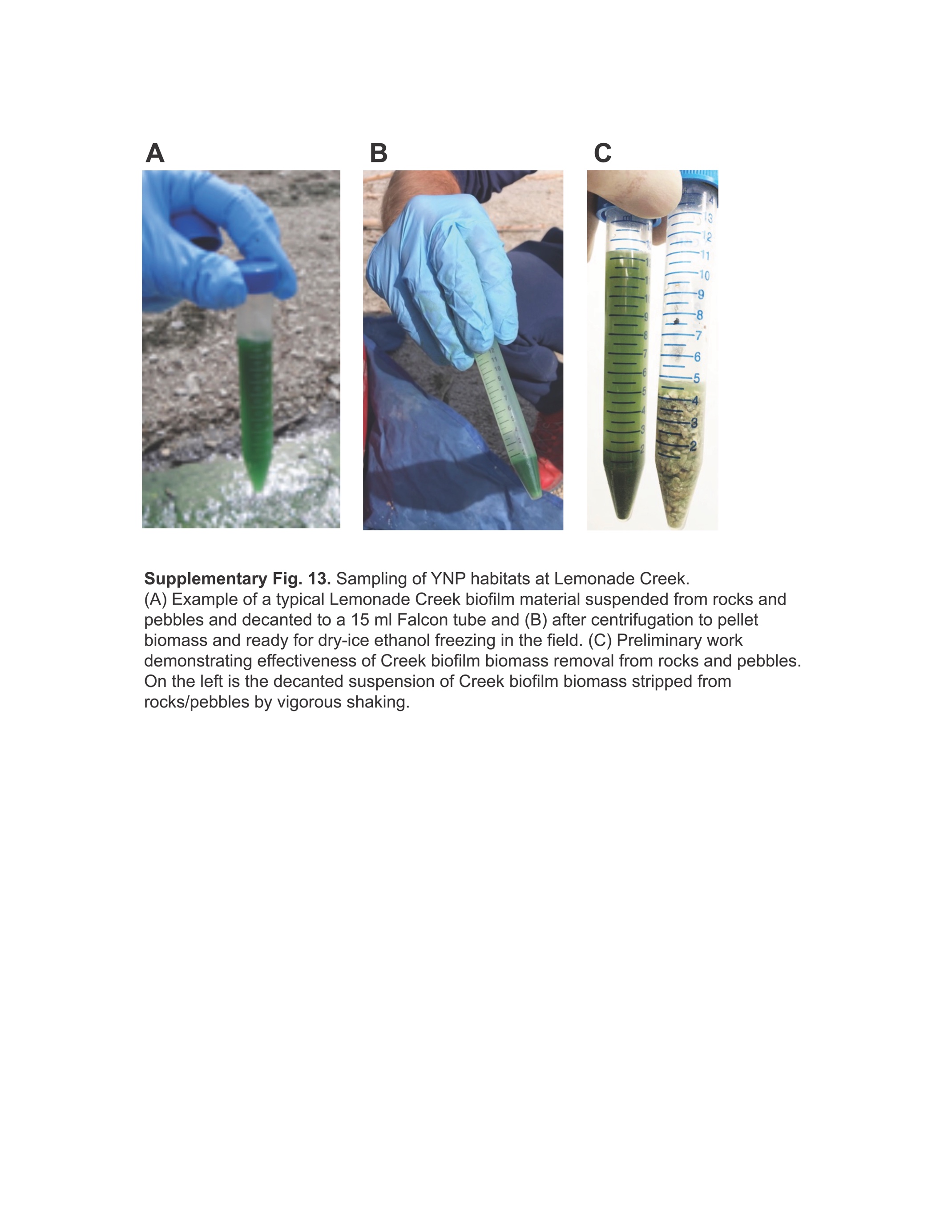

**Fig. S13. Sampling of YNP habitats at Lemonade Creek.** (A) Example of a typical Lemonade Creek biofilm material suspended from rocks and pebbles and decanted to a 15 ml Falcon tube and (B) after centrifugation to pellet biomass and ready for dry-ice ethanol freezing in the field. (C) Preliminary work demonstrating effectiveness of Creek biofilm biomass removal from rocks and pebbles. On the left is the decanted suspension of Creek biofilm biomass stripped from rocks/pebbles by vigorous shaking.

**Table S1. (separate file)**

Proportion of reads mapped to the contigs from each MAG in each metagenome sample, excluding viral MAGs with only single contigs.

**Table S2. (separate file)**

Proportion of reads mapped to transcripts from each MAG and domain level classification in each Sample of the Poly-A selected libraries.

**Table S3. (separate file)**

Proportion of reads mapped to transcripts from each MAG in each Sample of the RiboMinus selected libraries.

**Table S4. (separate file)**

Number of differentially accumulated metabolites between each time point in each combination of environment-extraction samples.

**Table S5. (separate file)**

Amount of metagenome reads generated from each sample before and after filtering and base correction.

**Table S6. (separate file)**

Statistics for the metagenome assembly generated from each sample.

**Table S7. (separate file)**

Rates of read mapping for all pair-wise combinations of metagenomes.

**Table S8. (separate file)**

Cyanidiophyceae genomes used for comparative analysis with the assembled metagenomes.

**Table S9. (separate file)**

Assembly quality and completeness statistics for extracted Cyanidiophyceae metagenome scaffolds and reference genomes.

**Table S10. (separate file)**

Average Nucleotide Identity (ANI) between the extracted cyanidiophyceae metagenome scaffolds and reference genomes.

**Table S11. (separate file)**

Quality and completeness of the sample-specific prokaryotic MAGs before and after removal of putative contaminant contigs by MAGpurify, MDMcleaner, and whokaryote.

**Table S12. (separate file)**

Statistics for the metagenome co-assembly generated from environment from the reads of unbinned contigs.

**Table S13. (separate file)**

Quality, completeness, and taxonomy of the final eukaryotic MAGs.

**Table S14. (separate file)**

Quality, completeness, and taxonomy of the final non-redundant prokaryotic MAGs.

**Table S15. (separate file)**

Amount of metatranscriptome reads generated from each sample before and after filtering and base correction.

**Table S16. (separate file)**

Rates of read mapping for each metatranscriptome assembly against the CDS of the predicted genes from a database of non-redundant MAG and Cyanidiophyceae reference genomes.

**Table S17. (separate file)**

Number of significantly differentially expressed genes between each pair-wise combination of time point within each combination of environment and Library method.

**Table S18. (separate file)**

Number genes from the arsenic and mercury detoxification pathways in each target orthogroup from each examined inflation parameter.

**Table S19. (separate file)**

Genes (functional annotations shown where available) in each target orthogroup from each examined inflation parameter.

**Table S20. (separate file)**

OG terms enriched in genes from the top 5% SNP block.

**Table S21. (separate file)**

List of *C. merolae* and *G. yellowstonensis* genes annotated as being part of cell cycle control pathways.
